# The PIWI protein Aubergine recruits eIF3 to activate translation in the germ plasm

**DOI:** 10.1101/859561

**Authors:** Anne Ramat, Maria-Rosa Garcia-Silva, Camille Jahan, Rima Naït-Saïdi, Jérémy Dufourt, Céline Garret, Aymeric Chartier, Julie Cremaschi, Vipul Patel, Mathilde Decourcelle, Amandine Bastide, François Juge, Martine Simonelig

## Abstract

Piwi-interacting RNAs (piRNAs) and PIWI proteins are essential in germ cells to repress transposons and regulate mRNAs. In *Drosophila*, piRNAs bound to the PIWI protein Aubergine (Aub) are transferred maternally to the embryo and regulate maternal mRNA stability through two opposite roles. They target mRNAs by incomplete base-pairing, leading to both their destabilization in the soma, and stabilization in the germ plasm. Here, we report a function of Aub in translation. Aub is required for translational activation of *nanos* mRNA, a key determinant of the germ plasm. Aub physically interacts with the poly(A) binding protein PABP and the translation initiation factor eIF3. Polysome gradient profiling identifies Aub role at the initiation step of translation. In the germ plasm, PABP and eIF3d assemble in foci that surround Aub-containing germ granules, and Aub acts with eIF3d to promote *nanos* translation. These results reveal a new mode of mRNA regulation by Aub, highlighting PIWI protein versatility in mRNA regulation.

## Introduction

Translational control is a widespread mechanism to regulate gene expression in many biological contexts. This regulation has an essential role during early embryogenesis, before transcription of the zygotic genome has actually started. In *Drosophila*, embryonic patterning depends on the translational control of a small number of maternal mRNAs (*1*). Among them, *nanos* (*nos*) mRNA encodes a key posterior determinant required for abdominal segmentation and development of the germline (*2*). *nos* mRNA is present in the whole embryo, but a small proportion accumulates at the posterior pole in the germ plasm, a specialized cytoplasm in which the germline develops (*3, 4*). Localization and translational control of *nos* mRNA are linked, such that the pool of *nos* mRNA present in the bulk of the embryo is translationaly repressed, whereas the pool of *nos* mRNA localized in the germ plasm is translationaly activated to produce a Nos protein gradient from the posterior pole (*3, 5, 6*). Both repression of *nos* mRNA translation in the bulk of the embryo and activation in the germ plasm are required for embryonic development.

The coupling between mRNA localization and translational control depends in part on the implication of the same factors in both processes. The Smaug (Smg) RNA binding protein specifically recognizes *nos* mRNA through binding to two Smaug recognition elements (SRE) in its 3’UTR (*7, 8*). Smg is both a translational repressor of *nos*, and a localization factor through its role in mRNA deadenylation and decay in the bulk of the embryo, by the recruitment of the CCR4-NOT deadenylation complex (*7–9*). Smg directly interacts with the Oskar (Osk) protein that is specifically synthetized at the posterior pole of oocytes and embryos and drives germ plasm assembly (*7, 10*). Smg interaction with Osk prevents Smg binding to *nos* mRNA, thus contributing to relieve both Smg-dependent translational repression and mRNA decay in the germ plasm (*7, 9, 11*). Osk is therefore a key player in the switch of *nos* and other germ cell mRNA regulation between soma and germ plasm of the embryo.

More recently, we have demonstrated the role of Aubergine (Aub) in the localization germ cell mRNAs to the germ plasm (*12, 13*). Aub is one of the three PIWI proteins in *Drosophila*. PIWI proteins belong to a specific clade of Argonaute proteins that bind 23-30 nucleotides (nt)-long small RNAs referred to as Piwi-interacting RNAs (piRNAs) (*14, 15*). piRNAs and PIWI proteins have an established role in the repression of transposable elements in the germline of animals. piRNAs target transposable element mRNAs through complementarity and guide interaction with PIWI proteins that, in turn, cleave targeted mRNAs through their endonucleolytic activity. In addition to this role, piRNAs have a conserved function in the regulation of cellular mRNAs in various biological contexts (*16*). In the *Drosophila* embryo, Aub loaded with piRNAs produced in the female germline is present both at low levels in the bulk of the embryo, and at higher levels in the germ plasm (*17, 18*). Aub binds maternal germ cell mRNAs through incomplete base-pairing with piRNAs (*12, 18, 19*). Aub binding to these mRNAs induce their decay in the bulk of the embryo, either by direct cleavage, or the recruitment together with Smg of the CCR4-NOT deadenylation complex (*12, 18*). In contrast, in the germ plasm Aub recruits Wispy, the germline-specific cytoplasmic poly(A) polymerase, leading to poly(A) tail elongation and stabilization of Aub-bound mRNAs (*13*). Thus, Aub and piRNAs play a central role in the localization of germ cell mRNAs through two opposite functions in mRNA stability: mRNA destabilization in the bulk of the embryo and stabilization in the germ plasm. The role of piRNAs and PIWI proteins in cellular mRNA regulation in other contexts, including mouse spermiogenesis and sex determination in *Bombyx* also depends on their function in the regulation of mRNA stability (*20–24*).

Here, we describe translational activation as a new mechanism of mRNA regulation by piRNAs and PIWI proteins. Using ectopic expression of Osk in the whole embryo to mimic the germ plasm, we show that Aub and piRNAs are required for *nos* mRNA translation. Mass spectrometry analysis of Aub interactors in early embryos identifies several components of the translation machinery, including translation initiation factors. We find that Aub physically interacts with the poly(A) binding protein (PABP) and several subunits of the translation initiation complex eIF3. Furthermore, PABP and eIF3d accumulate in foci that assemble around and partially overlap Aub-containing germ granules in the germ plasm. Polysome gradient analysis reveals that Aub activates translation at the initiation step. Finally, functional experiments involving the concomitant decrease of Aub and eIF3d show that both proteins act together in *nos* mRNA translation in the germ plasm. These results identify translational activation as a new level of mRNA regulation by PIWI proteins. Moreover, they expand the role of the general eIF3 translation initiation complex in translation regulatory mechanisms required for developmental processes.

## Results

### Aub is required for *nos* mRNA translation

Only a low amount (4%) of *nos* mRNA is localized to the germ plasm and actually translated (*3, 4*). *nos* mRNA stabilization and translation in the germ plasm depend on the presence of Osk. Therefore, we ectopically expressed Osk in the whole embryo using *UASp-osk* (*25*) and the germline-specific driver *nos-Gal4*, in order to increase translated *nos* mRNA levels and address the mechanisms of translational activation. Osk overexpression (*osk-OE*) in embryos from *UASp-osk/+; nos-Gal4/+* females led to increased and ectopic Nos protein synthesis in whole embryo (Fig. 1A). Quantification of Nos protein levels in *osk-OE* embryos, either following Nos visualization using immunostaining, or western blots, revealed a two-fold increase compared to wild-type embryos (Fig. 1A, B). In contrast, *nos* mRNA levels quantified using RT-qPCR were similar in *osk-OE* and wild-type embryos (Fig. 1C). This is consistent with the presence of high amounts of *nos* mRNA in the bulk of embryos, and *nos* spatial regulation depending mostly on translational control at these stages (0-2 hour-embryos). Therefore, Osk overexpression in 0-2 hour-embryos led to ectopic translational activation of *nos* mRNA, without changes in *nos* mRNA levels.

**Figure 1.**
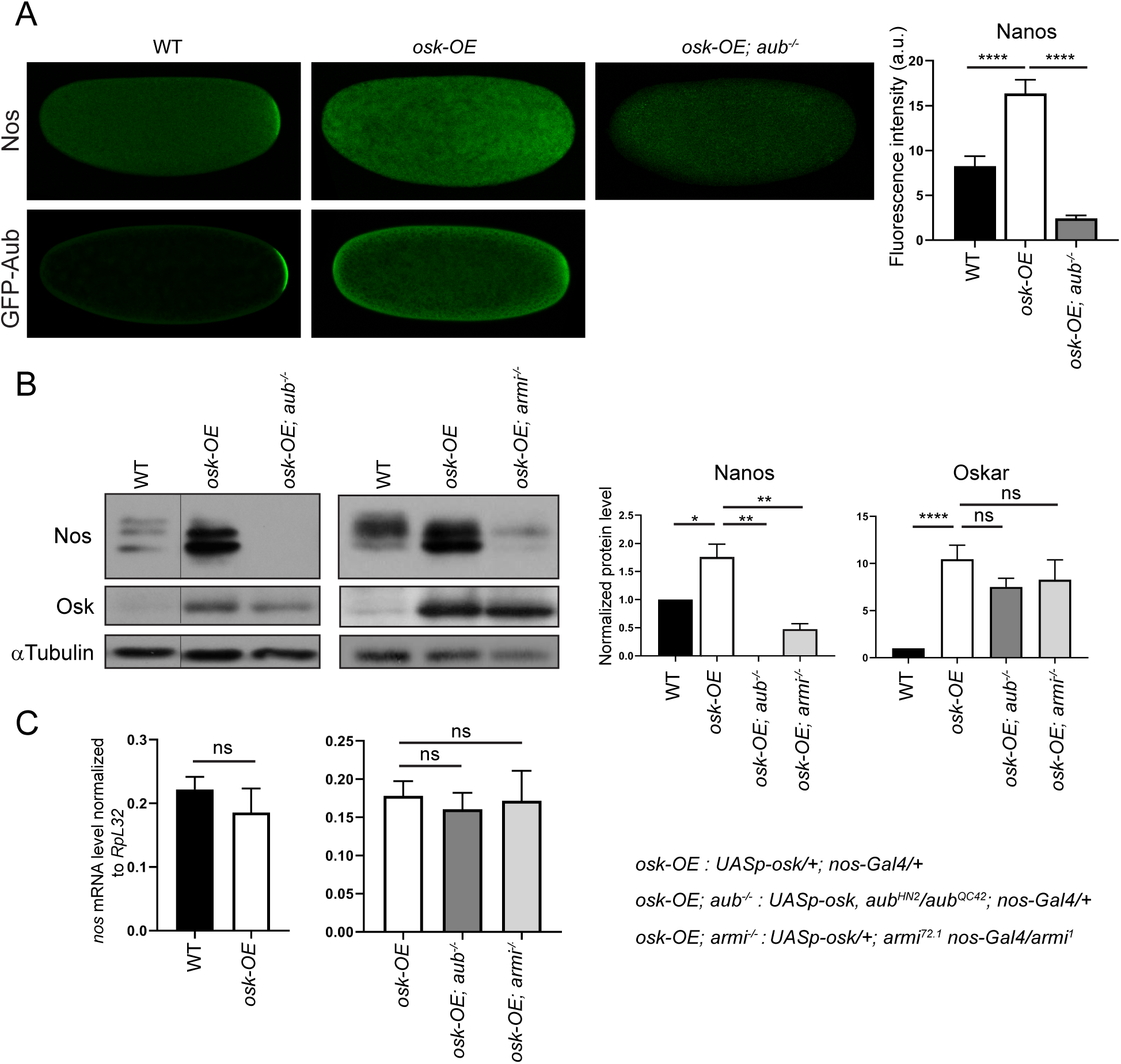
Aub and Armi are required for *nos* mRNA translation. (**A**) Immunostaining of wild-type (WT), *osk-OE* and *osk-OE; aub^-/-^* embryos with anti-Nos antibody (Top panels). The genotypes are indicated in the Figure. Fluorescence quantification was performed using the ImageJ software with 5 to 6 embryos per genotype. Error bars represent sem. **** *p*-value<0.0001 using the unpaired Student’s *t*-test. Immunostaining of *UASp-GFP-Aub nos-Gal4* and *UASp-osk/+; UASp-GFP-Aub nos-Gal4/+* embryos with anti-GFP antibody, showing the distribution of Aub protein (Bottom panels). Posterior is to the right. (**B**) Western blots of wild-type (WT), *osk-OE*, *osk-OE; aub^-/-^* and *osk-OE; armi^-/-^* embryos revealed with anti-Nos, anti-Osk and anti-αTubulin antibodies. αTubulin was used as a loading control. Quantification was performed using the ImageJ software with 3 to 6 biological replicates. Error bars represent sem. **** *p*-value<0.0001, \*\**p*-value<0.01, \**p*-value<0.05, ns: non significant, using the unpaired Student’s *t*-test. (**C**) Quantification of *nos* mRNA using RT-qPCR in wild-type (WT), *osk-OE*, *osk-OE; aub^-/-^* and *osk-OE; armi^-/-^* embryos. mRNA levels were normalized with *RpL32* mRNA. Quantification of 4 to 8 biological replicates. Error bars represent sem. ns: non significant, using the unpaired Student’s *t*-test.

Aub protein is present at low levels in the bulk of wild-type embryos and highly accumulates in the germ plasm (*18*). Ectopic expression of Osk led to an homogeneous redistribution of Aub in the embryo (Fig. 1A). Strikingly, the lack of Aub in *osk-OE* embryos resulted in the lack of Nos protein synthesis (Fig. 1A, B), although *nos* mRNA levels were not decreased (Fig. 1C). This result suggested that Aub was required for *nos* mRNA translational activation in the presence of Osk. Importantly, the level of Osk protein was not significantly affected by *aub* mutation, indicating that the lack of Nos protein did not result from the lack of Osk (Fig. 1B). Of note, the *UASp-osk* transgene almost exclusively overexpressed the long Osk isoform of the two isoforms, Short-Osk and Long-Osk (Fig. S1A). Long-Osk can induce germ plasm assembly when overexpressed -although less actively than Short-Osk-(*26*). Long-Osk levels were poorly affected by *aub* mutations, making this *UASp-osk* transgene a useful tool to address direct *nos* mRNA regulation by Aub and piRNAs (Fig. 1B, S1A).

We analyzed the role of Armitage (Armi), another component of the piRNA pathway with a prominent role in piRNA biogenesis (*27*), on *nos* mRNA translational activation. Nos protein levels were strongly reduced in *osk-OE; armi^-/-^* embryos, compared to *osk-OE* embryos, although again, the levels of Osk protein were not significantly decreased (Fig. 1B). *nos* mRNA levels quantified using RT-qPCR remained unaffected by *armi* mutation (Fig. 1C), revealing a role of Armi in *nos* mRNA translational control. Armi does not localize to the germ plasm (*13*). Instead the defect in *nos* mRNA translational activation in *armi* mutant might depend on highly reduced piRNA levels in this mutant (*27*), suggesting that piRNAs were required for Aub binding to *nos* mRNA for its role in translational activation. This is consistent with Aub iCLIP assays showing that an Aub double point mutant in the PAZ domain, Aub^AA^ that is unable to load piRNAs, was also unable to bind mRNAs (*12*).

These data are consistent with a direct role of Aub and piRNAs in *nos* mRNA translation through piRNA-dependent binding of *nos* by Aub. In this hypothesis, a piRNA pathway component specifically involved in transposable element regulation should not interfere with *nos* mRNA translational control. We used Panoramix (Panx), a key factor in Piwi-dependent transcriptional silencing of transposable elements, which acts downstream of Piwi and has no function in piRNA biogenesis (*28, 29*). *panx* mutants had no effect on Nos protein levels in *osk-OE* embryos, consistent with a role of Aub and piRNAs in *nos* mRNA translation, independent of their role in transposable element regulation (Fig. S1B).

Finally, most *aub* mutant embryos fail to develop, although they are fertilized (*12, 30*). To address whether the lack of Nos protein in *osk-OE; aub^-/-^* embryos could result from their arrest of embryonic development, we quantified Nos protein levels in *osk-OE* unfertilized eggs that are activated by egg laying but do not develop. Nos levels were similar in *osk-OE* unfertilized eggs and embryos demonstrating that the defect in Nos protein synthesis in *osk-OE;* aub^-/-^ embryos did not result from their lack of embryonic development (Fig. S1C).

Together, these results show that Aub binding to *nos* mRNA plays a direct role in translational activation in the presence of Osk.

### Ectopic expression of Osk leads to the formation of germ granule-like granules in the soma

In the germ plasm, Osk leads to the assembly of germ granules that are large ribonucleoprotein particles containing mRNAs required for germ cell specification and development (*4, 31*). In addition to Osk, Aub is a core component of germ granules (*32*). We asked whether, Osk ectopic expression in the somatic part of the embryo could lead to the formation of RNA granules related to germ granules, containing Aub and *nos* mRNA. Immunostaining of *osk-OE* embryos also expressing GFP-Aub revealed that Osk was present in the bulk of the embryo where it accumulated in cytoplasmic foci that became larger around nuclei (Fig. 2A). GFP-Aub was also present in cytoplasmic foci in the bulk of *osk-OE* embryos and in larger foci around nuclei. Small foci of either Osk or GFP-Aub were dispersed in the cytoplasm and did not colocalize. In contrast, Osk and GFP-Aub colocalized in larger foci that surrounded nuclei, indicating a different composition of these large foci (Fig. 2A-A’’, D). Single molecule FISH (smFISH) of *nos* mRNA in embryos of the same genotype showed that *nos* mRNA accumulated in larger foci around nuclei where it colocalized with GFP-Aub (Fig. 2B-B’’, D). Strikingly, Nos protein also accumulated around nuclei and partially colocalized with GFP-Aub in large foci, suggesting that *nos* mRNA translation occurred in the vicinity of these granules (Fig. 2C-C’’, D).

**Figure 2.**
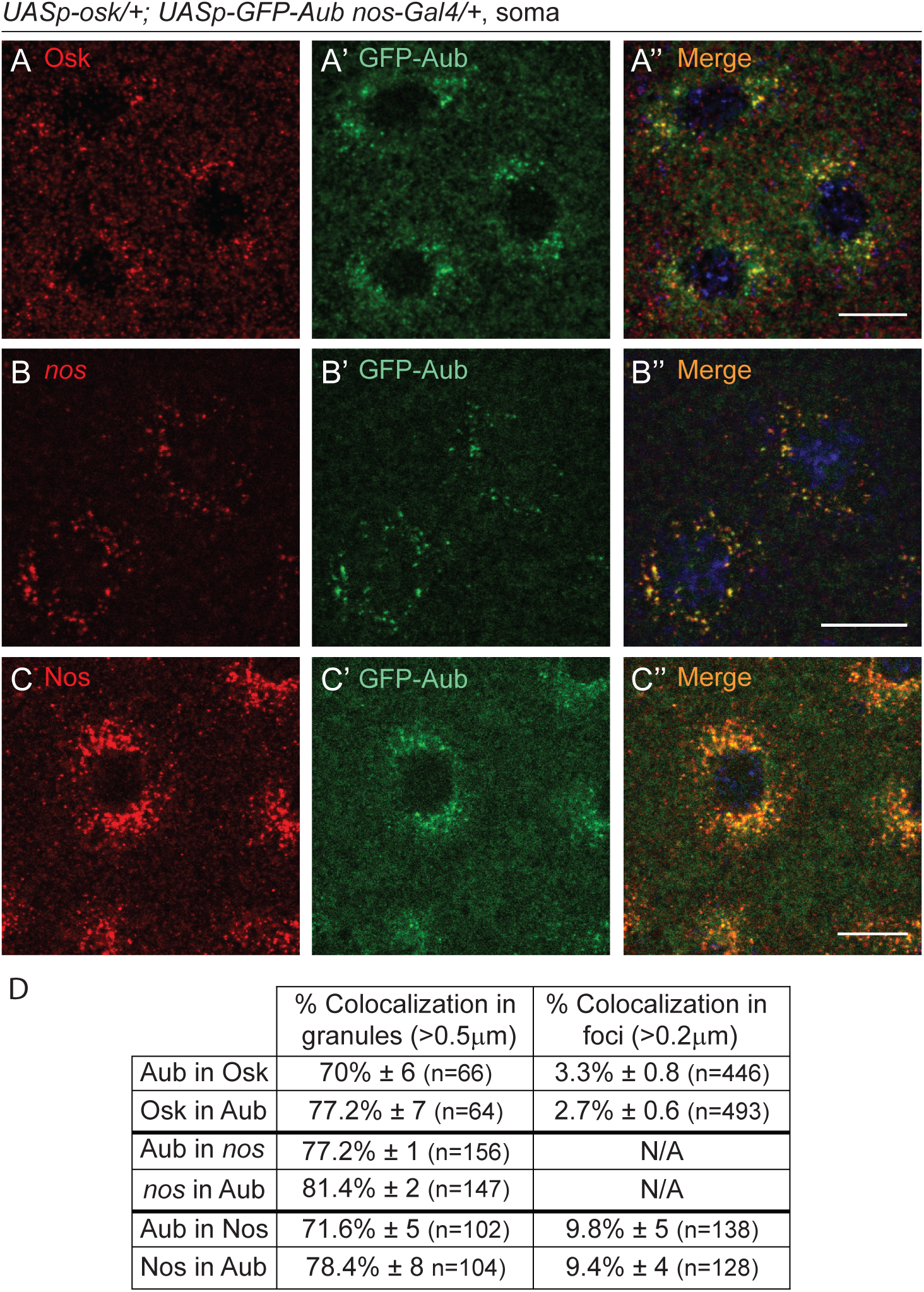
Ectopic expression of Osk nucleates RNA granules related to germ granules in the soma. (**A-C’’**) Immunostaining of *UASp-osk/+; UASp-GFP-Aub nos-Gal4/+* embryos with anti-Osk (red) and anti-GFP (green) to visualize Aub (**A-A’’**); and anti-Nos (red) and anti-GFP(green) (**C-C’’**); and smFISH of embryos with the same genotype revealing *nos* mRNA and GFP-Aub through GFP fluorescence (**B, B’’**). DNA was visualized using DAPI. (**D**) Quantification of colocalization of immunostaining and smFISH shown in **A-C’’**, using the Imaris software. Granules (>0.5 μm) and foci (>0.2 μm) were quantified around nuclei and in the cytoplasm between nuclei, respectively.

We conclude that the presence of Osk in the somatic part of *osk-OE* embryos induce the formation of RNA granules that share functional similarities with germ granules, in which Aub and *nos* mRNA accumulate and at the proximity of which *nos* mRNA is translated.

### Aub interacts with translation initiation factors

To further decipher the function of Aub, we identified Aub interactors in embryos. GFP-Aub was immunoprecipitated from *UASp-GFP-Aub nos-Gal4* 0-2 hour-embryos and the coprecipitated proteins were analyzed using mass spectrometry. Embryos expressing GFP alone were used as negative controls (Fig. S2A). 107 proteins were significantly enriched in GFP-Aub immunoprecipitation (IP) (*p*-value<0.05) (Table S1). Known Aub interactors were identified, including Tudor (Tud) that is restricted to the germ plasm and required for Aub accumulation in the germ plasm (*33, 34*), three components of the *nos* translation repressor complex, Trailer hitch (Tral), Belle (Bel) and Cup (*35*), and Capsuleen/PRMT5 (Csul) the methyltransferase responsible for Aub arginine dimethylation (*36*) (Fig. 3A). Several RNA binding proteins were also found in GFP-Aub IP (Fig. 3A). Importantly, six translation initiation factors were identified as Aub interactors, among which the poly(**A**) binding protein (PABP), three subunits of eIF3 (eIF3d, eIF3k and eIF3b), and eIF4E, another component of *nos* translation repressor complex (*35*) (Fig. 3B). In addition, 48 ribosomal proteins coprecipitated with Aub (Fig. S2B). Gene Ontology (GO) term enrichment analysis using FlyMine (http://www.flymine.org) identified “Translation” as the most enriched term among Aub interactors (Fig. 3E). We also analyzed Aub interactors in *osk^54^* mutant embryos that do not form germ plasm, with the aim of identifying specific Aub interactors in the germ plasm, which might be lost in *osk* mutant embryos. However, mass spectrometry of GFP-Aub IP from *osk^54^* mutant embryos identified a very similar set of proteins to that identified in *osk^+^* embryos (Fig. 3C, D, S2C, Table S1). These data suggested that Osk might not affect Aub interaction with its protein interactors, but rather activity of the Aub complex.

**Figure 3:**
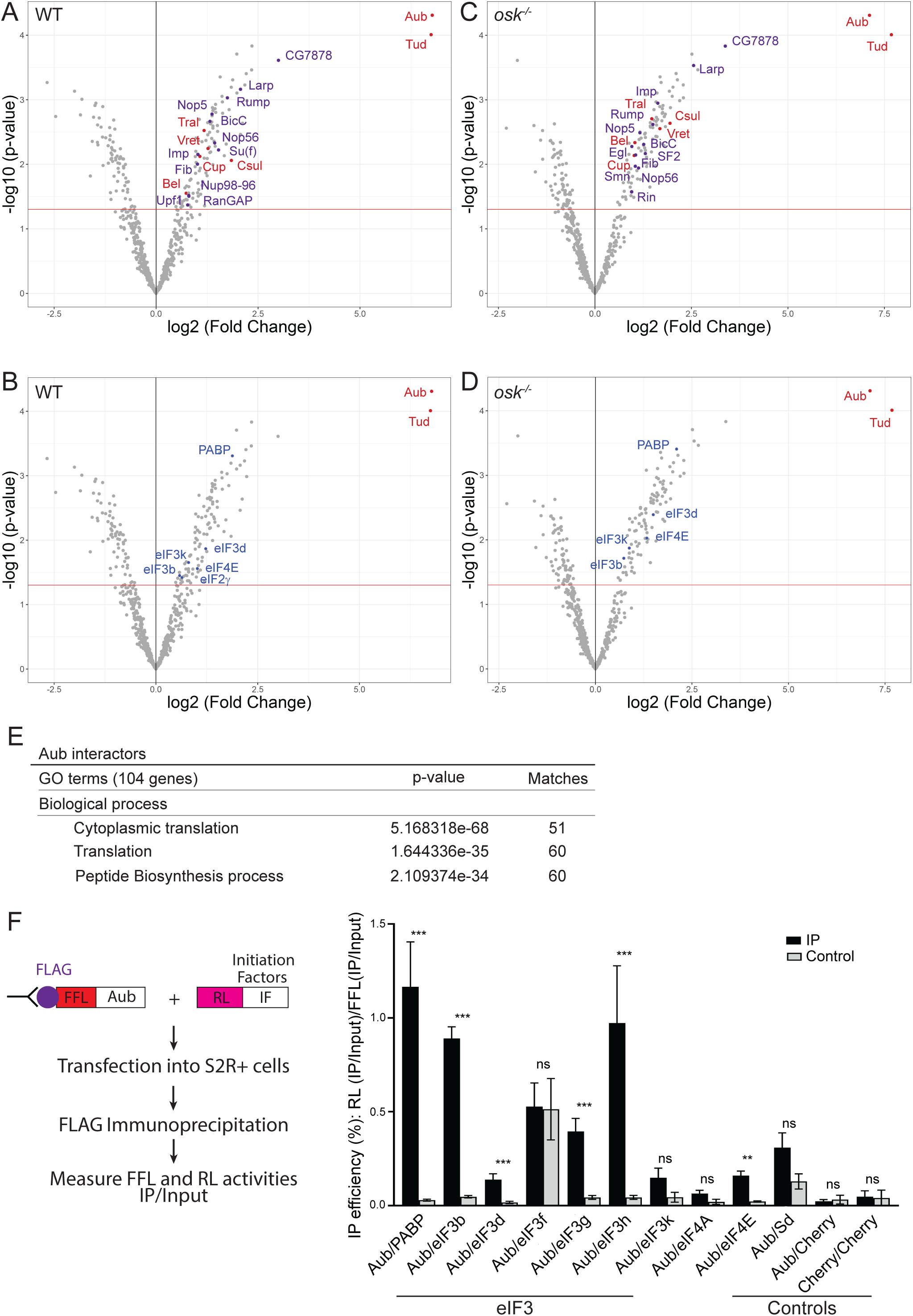
Identification of Aub interacting partners. (**A-D**) Volcano plots showing the mass spectrometry analysis of GFP-Aub immunoprecipitation from 0-2 hour-embryos. Embryos expressing cytoplasmic GFP were used as control. (**A, B**) *UASp-GFP-Aub nos-Gal4* embryos; (**C, D**) *osk^54^; UASp-GFP-Aub/nos-Gal4* embryos. The analysis was based on four biological replicates. The red line indicates the significance threshold (*p*-value = 0.05). Known Aub interactors and RNA binding proteins are indicated in red and purple, respectively (**A, C**); translation initiation factors are indicated in blue (**B, D**). (**E**) Gene ontology (GO) analysis of proteins identified as Aub interactors by mass spectrometry. (**F**) Validation of Aub interactors using the LUMIER assay. Left: schematic representation of the assay (FFL, Firefly luciferase; RL, Renilla luciferase). Right: graph plotting the IP efficiency of the indicated proteins. The values are IP efficiencies of the coprecipitation of the RL fusion proteins (IP/Input) normalized by the IP/Input values for FLAG-FFL-Aub. Error bars represent sd. Stars indicate values significantly greater than six-times the mean value obtained in the control IPs without anti-FLAG antibody (Control). Scalloped (Sd) and Cherry proteins were used as negative controls. *** *p*-value<0.001, ** *p*-value<0.01, ns: non significant, using the Z-test.

We used quantitative luminescence-based coIP (LUMIER) assays to validate Aub interactions with translation initiation factors (*37*). Aub was fused to FLAG-tagged Firefly luciferase, whereas potential interactors were fused to Renilla luciferase. Following transient expression in *Drosophila* S2R+ cells, Aub was immunoprecipitated with anti-FLAG antibodies, or without antibodies as negative control, and interactor coIP was quantified by recording Renilla and Firefly luciferase activities (Fig. 3F). PABP, four subunits of eIF3 (eIF3b, eIF3d, eIF3g and eIF3h) among six tested subunits, and eIF4E were found to significantly coprecipitate with Aub in these assays. Thus, although eIF3k that was identified as an Aub interactor by mass spectrometry could not be confirmed with the LUMIER assay, eIF3d and eIF3b interaction with Aub was confirmed, and two other eIF3 subunits, eIF3g and eIF3h were found in complex with Aub.

These results reveal that Aub physically interacts with the translation machinery and are consistent with a direct function of Aub in translation regulation.

### Aub interaction with PABP and eIF3d

Because PABP and eIF3d showed the strongest association with Aub in the mass spectrometry analysis, and have key roles in translation initiation, we further investigated their interaction with Aub. We used coIP to address Aub physical interaction with PABP in embryos. PABP coprecipitated with GFP-Aub in 0-2 hour-embryos; however, this coprecipitation was strongly reduced in the presence of RNase (Fig. 4A). In the reverse experiment, PABP was also able to coprecipitate Aub, but this coprecipitation was abolished in the presence of RNase (Fig. 4B). These results could indicate either that Aub and PABP did not interact directly and coprecipitated through their binding to the same mRNAs, or that Aub direct interaction with PABP was stabilized by mRNAs in a tripartite association. To address this question, we analyzed direct interaction between Aub and PABP using GST pull-down assays. Aub has three domains characteristic of Argonaute proteins (PAZ, MID and PIWI) and was separated into two parts, Aub(1–482) that contains the N-terminal and PAZ domains, and Aub(476–866) that contains the MID and PIWI domains (Fig. 4C). PABP is composed of four RNA recognition motifs (RRM1-4), a proline rich linker region and a PABC (PABP C-terminal) domain (*38*). Each RRM and the PABC domain were fused separately to GST. In vitro-translated HA-tagged Aub(1–482) bound to recombinant GST-RRM1, but not to the other PABP domains fused to GST or GST alone. In addition, HA-Aub(476–866) did not bind to any PABP domain (Fig. 4C). These data revealed direct interaction between RRM1 of PABP and the N-terminal half of Aub.

**Figure 4:**
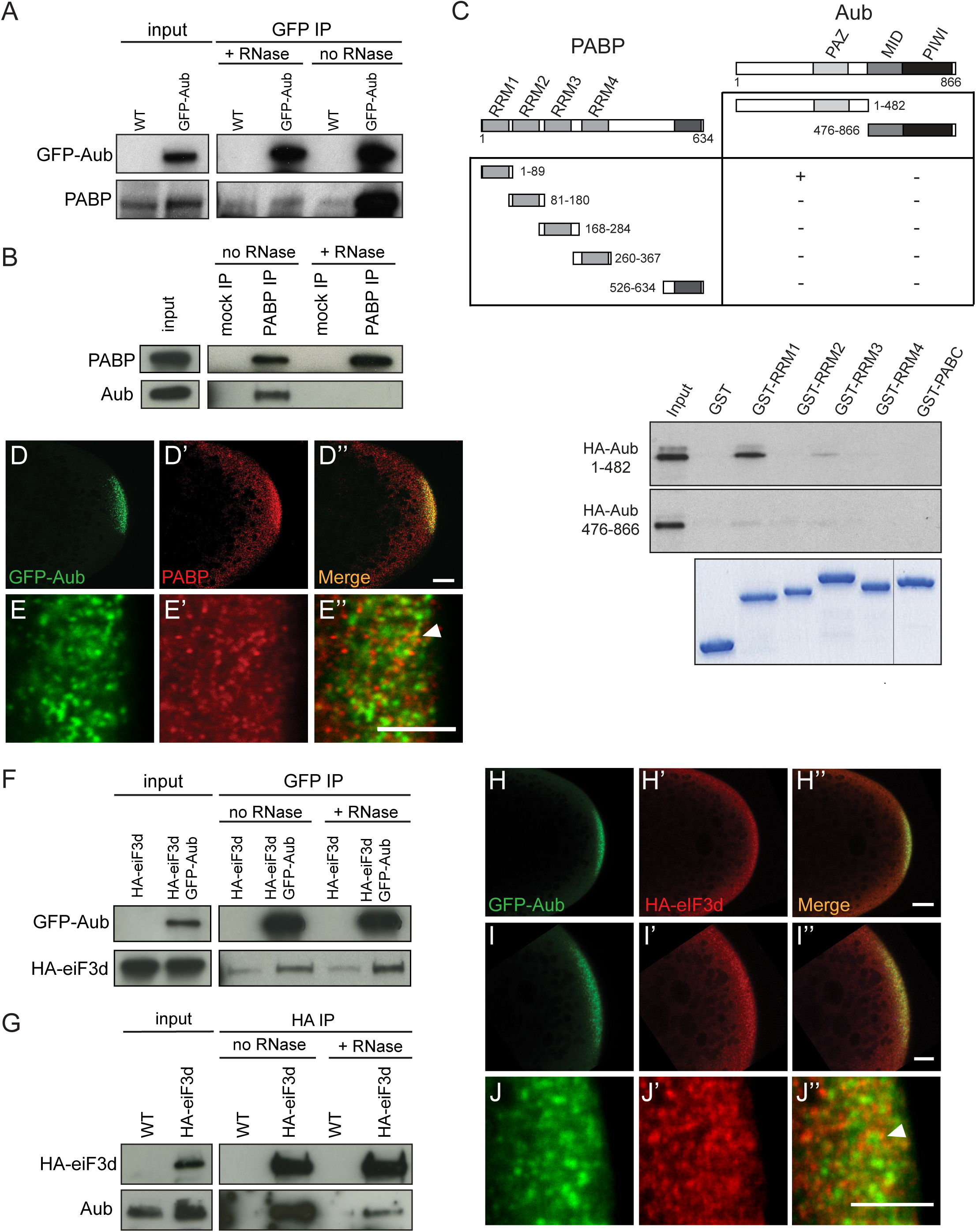
Aub physical interaction with PABP and eIF3d. (**A**) CoIP of PABP with GFP-Aub in 0-2 hour-embryos. Wild-type (WT, mock IP) or *nos-Gal4 UASp-GFP-Aub* (GFP IP) embryo extracts were immunoprecipitated with anti-GFP, either in the absence or the presence of RNase A. Western blots were revealed with anti-GFP and anti-PABP. Inputs are extracts before IP. (**B**) CoIP of Aub with PABP in 0-2 hour-embryos. Wild-type embryo extracts were immunoprecipitated with anti-PABP (PABP IP) or rabbit serum (mock IP), either in the absence or the presence of RNase A. Western blots were revealed with anti-PABP and anti-Aub. Inputs are extracts before IP. (**C**) GST pull-down assays between GST-PABP and HA-Aub. Constructs and interactions are shown in the table. HA-tagged Aub fragments were revealed using western blots with anti-HA. Inputs correspond to 1/10 of in vitro-synthetized HA-Aub fragments before pull-down. GST alone was used as negative control. GST and GST-recombinant proteins used in each pull down are shown in the bottom gel. (**D-E’’**) Immunostaining of *UASp-GFP-Aub nos-Gal4* embryos with anti-GFP (green) to visualize Aub and anti-PABP (red). Posterior of embryos are shown. Higher magnification showing the distribution of Aub-containing germ granules and PABP foci (**E-E’’**). Colocalization and overlap between Aub and PABP staining are quantified in Fig. S3A. The white arrowhead shows PABP foci surrounding a germ granule. Scale bars: 20 μm in **D** and 5 μm in **E**. (**F, G**) CoIP of HA-eIF3d with GFP-Aub (**F**) and of Aub with HA-eIF3d (**G**) in 0-2 hour-embryos. *UASp-HA-eIF3d/+; nos-Gal4/+* (mock IP) or *UASp-HA-eIF3d/+; UASp-GFP-Aub nos-Gal4/+* (GFP IP) embryo extracts were immunoprecipitated with anti-GFP, either in the absence or the presence of RNase A. Western blots were revealed with anti-GFP and anti-HA (**F**). Wild-type (WT, mock IP) or *UASp-HA-eIF3d/+; nos-Gal4/+* (HA IP) embryo extracts were immunoprecipitated with anti-HA, either in the absence or the presence of RNase A. Western blots were revealed with anti-HA and anti-Aub (**G**). Inputs are extracts before IP in **F** and **G**. (**H-J’’**) Immunostaining of *UASp-HA-eIF3d/+; UASp-GFP-Aub nos-Gal4/+* embryos with anti-GFP (green) to visualize Aub and anti-HA (red) to visualize eIF3d. Posterior of embryos are shown. Higher magnification showing the slight accumulation of HA-eIF3d at the posterior pole (**I-I’’**), and the distribution of Aub-containing germ granules and eIF3d foci (**J-J’’**). Colocalization and overlap between Aub and eIF3d staining are quantified in Fig. S3D. The white arrowhead shows eIF3d foci surrounding a germ granule. Scale bars: 20 μm in **H**, 10 μm in **I** and 5 μm in **J**.

We then analyzed potential colocalization of Aub and PABP. Co-immunostaining of GFP-Aub-expressing embryos with anti-PABP and anti-GFP antibodies showed that PABP was distributed in the whole embryo and specifically accumulated in the germ plasm (Fig. 4D-D’’). Strikingly, PABP was present in foci, and in the germ plasm most of these foci (81.3 to 88.7%, Fig. S3A) either colocalized or were in close proximity to and surrounded Aub-containing germ granules, with a partial overlap of both proteins (Fig. 4E-E’’, S3A).

eIF3 is composed of twelve subunits and one associated factor, and coordinates several steps of translation initiation (*39*). Interestingly, in addition to this role in basal translation, eIF3 plays regulatory roles in the translation of specific mRNAs. eIF3d appears to be a major actor in eIF3 regulatory functions, either through its binding of 5’UTR of specific mRNAs, leading to cap-independent translation, or directly through its interaction with the cap structure (*40, 41*). Aub interaction with eIF3d was analyzed in embryos using coIP. GFP-Aub was able to coprecipitate HA-eIF3d in 0-2 hour-embryos, and this coprecipitation was maintained in the presence of RNase (Fig. 4F). Conversely, HA-eIF3d was able to coprecipitate Aub in 0-2 hour-embryos, and although less efficient, this coprecipitation remained in the presence of RNase (Fig. 4G). Colocalization of Aub and eIF3d was analyzed in embryos expressing both GFP-Aub and HA-eIF3d. eIF3d was present in the whole embryo with a slight accumulation in the germ plasm (Fig. 4H-I’’, S3B, S3C). Similarly to PABP, eIF3d formed foci, a large number of which colocalized with or surrounded Aub-containing germ granules in the germ plasm (72.4 to 81.6%), with a partial colocalization of both proteins at the edge of the granules (Fig. 4J-J’’, S3D).

Taken together, these results show that Aub is in complex with the translation initiation factors PABP and eIF3d. In the germ plasm, PABP and eIF3d have a specific organization around germ granules and colocalize with Aub at the periphery of the granules, suggesting that translation might take place at the edge of germ granules.

### Mechanism of Aub-dependent translational activation

Aub association with translation initiation factors suggested that Aub might activate *nos* mRNA translation at the level of initiation. We directly addressed this question using polysome profiling in which mRNA-protein complexes are separated by fractionation through linear sucrose gradients (*42*). mRNA localization within the sucrose gradient reflects its translation status: migration in the light RNP or monosomal fractions of the gradient indicates a lack of translation, whereas migration in the heavy polysomal fractions indicates active translation. Polysome profiling was performed with 0-2 hour-wild-type, -*osk-OE* and *-osk-OE; aub^-/-^* embryos. The abundance of polysomes was reduced in *osk-OE* embryos compared to wild-type, indicating that ectopic expression of Osk in the whole embryo affected basal translation (Fig. 5A). In contrast, polysome abundance was partially restored in *osk-OE; aub^-/-^* embryos, revealing that translation was active in these embryos (Fig. 5A). Thus, the level of basal translation was affected oppositely to the level of Nos protein. This is consistent with Aub being involved in a regulatory mode of translation occurring on specific mRNAs. To confirm this point, we used *smg* mRNA as a control, since it is highly translated in the whole embryo upon egg activation (*43, 44*). Smg protein levels were not decreased in *osk-OE; aub^-/-^* embryos compared to *osk-OE* embryos, confirming the specificity of Aub-dependent translational activation (Fig. S4A). Western blot analysis of the gradient fractions revealed association of Aub with actively translating mRNAs in the heavy polysomal fractions, and the co-association of PABP in these fractions (Fig. 5E, F). To confirm Aub association with actively translating mRNAs, we treated embryo lysates with puromycin that causes premature termination of elongating ribosomes. Puromycin treatment efficiency was validated by the complete disassembly of polysomes visualized by absorbance measurement at OD 254 nm, and the shift of ribosomal proteins to monosomal and lighter fractions containing 60S and 40S ribosomal subunits (Fig. 5E, F). Aub shifted to the light mRNP fractions in the presence of puromycin indicating its *bona fide* association with translating mRNAs. In contrast, although PABP was shifted towards lighter fractions of the gradients in the presence of puromycin, a certain amount remained present in most fractions, suggesting the presence of heavy RNA complexes containing PABP in *Drosophila* embryos (Fig. 5F). This is consistent with the presence of mRNAs in heavy fractions of sucrose gradients independently of translation, in polysome gradients from early embryos (*45*). We then quantified mRNA through polysome gradients using RT-qPCR. *nos* mRNA was mostly present in initiation and light polysomal fractions in wild-type embryos, in agreement with a low amount of *nos* mRNA being actively translated (Fig. 5B). In *osk-OE* embryos, the level of *nos* mRNA decreased in the initiation fractions whereas it increased in the heavy polysomal fractions, consistent with the two-fold increase of Nos protein levels in these embryos (Fig. 5B, 1A, B).

**Figure 5.**
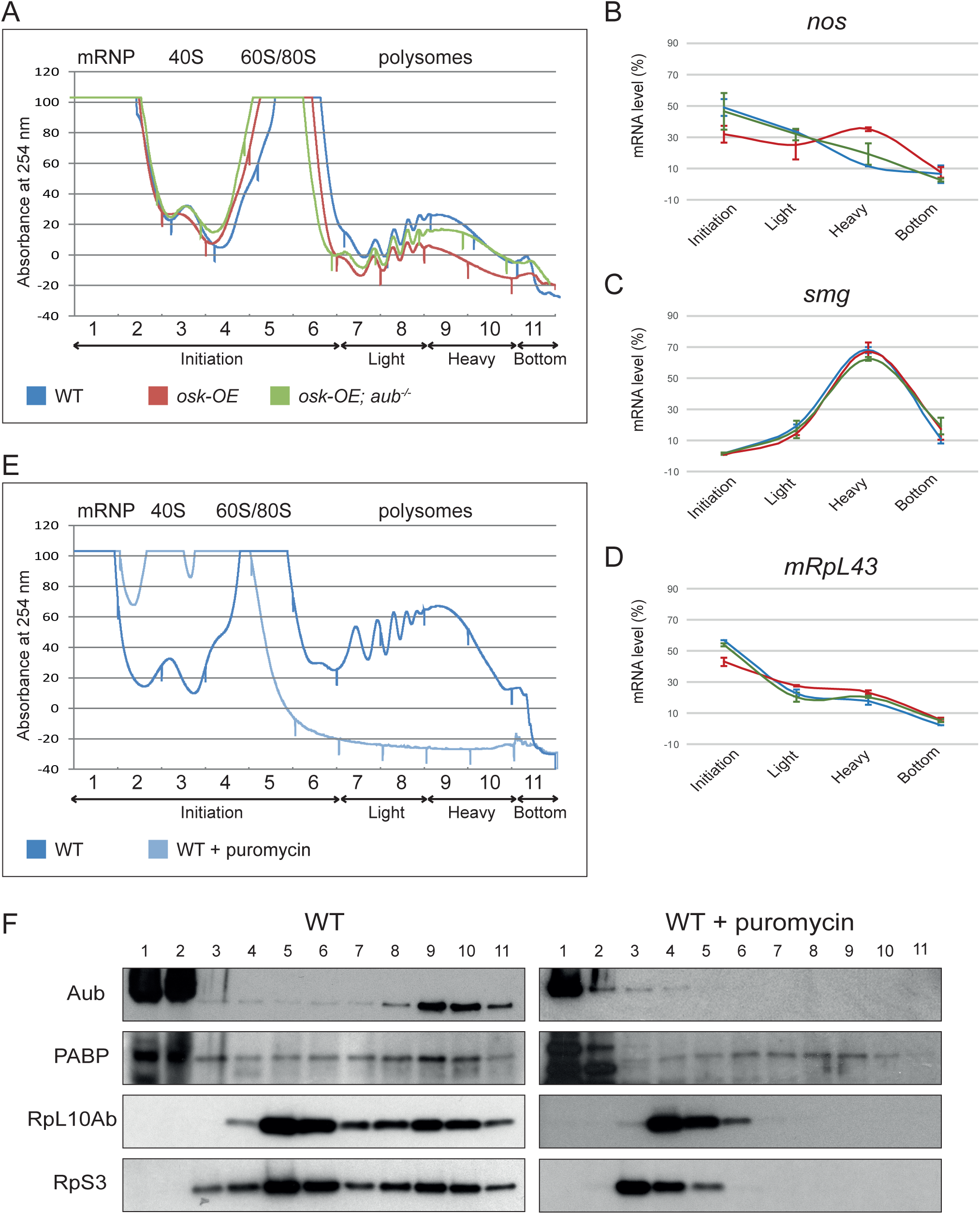
Aub acts at the level of translation initiation. (**A**) Profile of 254 nm absorbance for 0-2 hour-wild-type (WT, blue), -*osk-OE* (red) and -*osk-OE; aub^-/-^* (green) embryo extracts fractionated into 10-50 % sucrose gradients. Complete genotypes are as in Fig. 1. Fractions were pooled as indicated on the graph into: initiation (fractions 1 to 6), light polysomes (fractions 7 and 8), heavy polysomes (fractions 9 and 10) and bottom (fraction 11). (**B-D**) Quantification of *nos* (**A**), *smg* (**B**) and *mRpL43* (**C**) mRNAs using RT-qPCR in the different fractions of the gradients for wild-type- (WT, blue), -*osk-OE* (red) and -*osk-OE; aub-/-* (green) embryos. mRNA levels are indicated in percentage of total mRNA in all the fractions. Mean of two biological replicates, quantified in triplicates. Error bars represent sem. *smg* and *mRpL43* were used as control mRNAs. (**E**) Profile of 254 nm absorbance for 0-2 hour-wild-type (WT) embryos treated (light blue), or not (dark blue) with puromycin, fractionated into 10-50 % sucrose gradients. (**F**) Western blot showing the distribution of Aub and PABP through the gradient from wild-type (WT) embryos treated or not with puromycin. Two ribosomal proteins, RpL10Ab (60S ribosome subunit) and RpS3 (40S ribosome subunit) were used to record puromycin treatment efficacy.

Quantification of *nos* mRNA through the gradient in the presence of puromycin confirmed that the pool of *nos* present in the heavy fractions was indeed associated with actively translating polysomes (Fig. S4B-D). Interestingly, in *osk-OE; aub^-/-^* embryos, the distribution of *nos* mRNA was similar to that in wild-type embryos, with high amounts of mRNA in initiation fractions (Fig. 5A). These results revealed the role of Aub at the initiation step of translation. To further confirm the role of Aub in translational activation of specific mRNAs, we quantified *smg* and *mRpL43* mRNAs through the polysome gradients. Consistent with *smg* active translation in early embryos, most *smg* mRNA was present in heavy polysomal fractions, and this profile was not affected in *osk-OE* and *osk-OE; aub^-/-^* embryos, indicating that *smg* translation was independent of both Osk and Aub (Fig. 5C). *mRpL43* was used as a control mRNA that is not bound by Aub (*12*) and similarly, its distribution through the gradient was not strongly affected in *osk-OE* and *osk-OE; aub^-/-^* embryos (Fig. 5D).

These results show that Aub plays a role in the translation of specific mRNAs and acts at the level of translation initiation.

### eIF3d plays a role in Aub-dependent translational activation

To address the biological relevance of Aub/eIF3d physical interaction we analyzed the effect of the concomitant reduction of *aub* and *eIF3d* gene dosage by half. Although independent *aub* or *eiF3d* heterozygous mutant embryos showed a low level of lethality (2 to 3%), embryonic lethality significantly increased up to 21% in double heterozygous mutants, suggesting that Aub and eIF3d act together in embryonic development (Fig. 6A). *nos* mRNA translation was then recorded in these embryos using immunostaining. The Nos protein level visualized by immunofluorescence at the posterior cortex was quantified. In the wild-type, 86% of embryos showed a full accumulation of Nos protein at the posterior pole, whereas 14% had a reduced accumulation (Fig. 6B, C). Nos accumulation in heterozygous *aub* or *eIF3d* mutant embryos was similar to wild type. In contrast, in *aub^-/+^; eIF3d^-/+^* double heterozygous mutants, the percentage of embryos with reduced Nos accumulation significantly increased to 35% (Fig. 6B, C). This reduction of Nos accumulation did not correlate with reduced Osk accumulation or reduced *nos* mRNA levels at the posterior pole (Fig. 6C, S5), indicating a direct defect in *nos* mRNA translation. We conclude that Aub/eIF3d physical interaction is required for *nos* mRNA translational activation.

**Figure 6:**
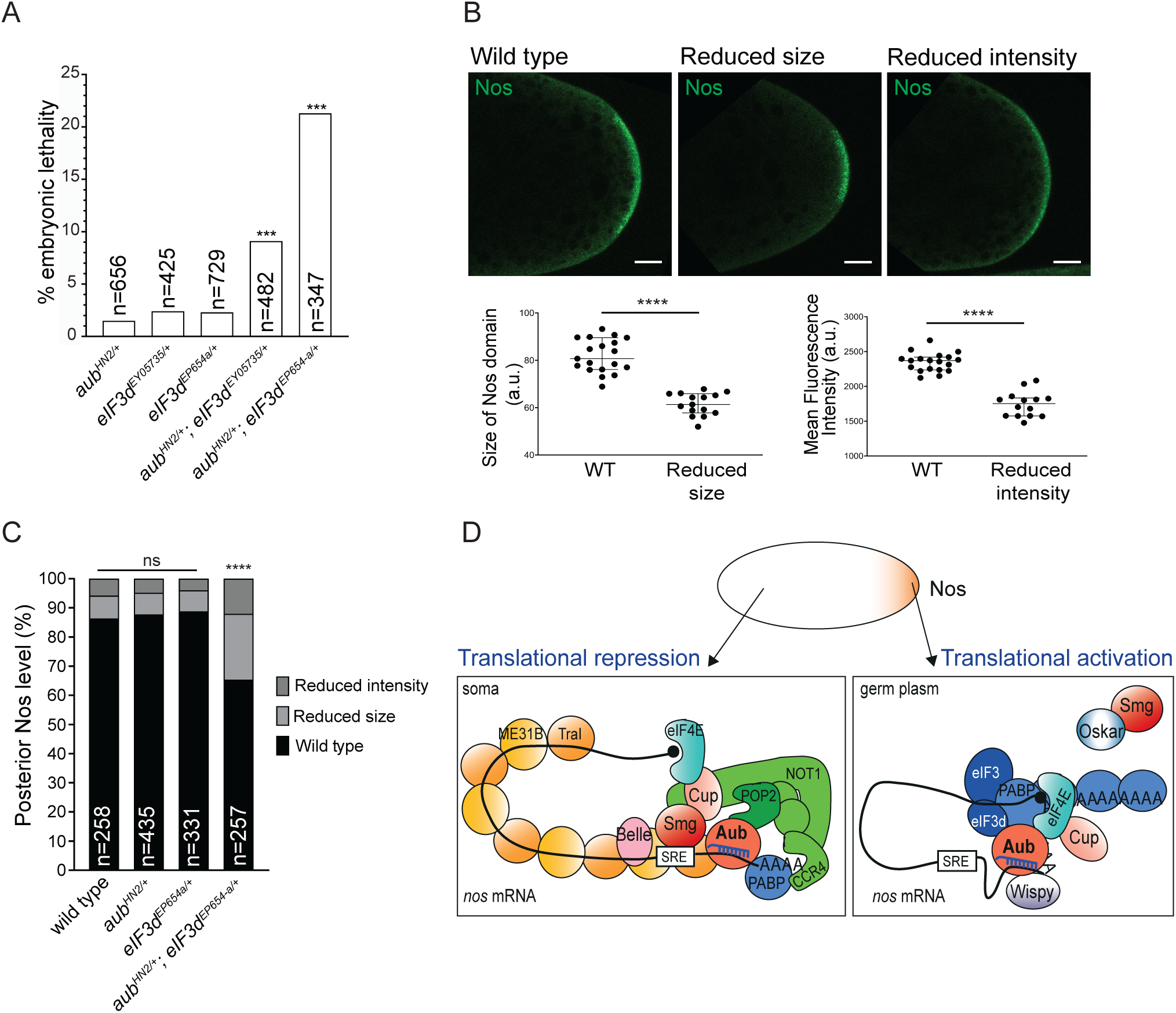
Aub and eIF3d functionally interact for *nos* mRNA translation. (**A**) Percentage of embryonic lethality of single or double *aub* and *eIF3d* heterozygous mutants. The genotypes are indicated. *** *p*-value<0.001, using the χ2 test. (**B, C**) Immunostaining of single and double *aub* and *eIF3d* heterozygous mutant embryos with anti-Nos antibody. Posterior of embryos showing the three types of obtained staining: wild type, reduced size or reduced intensity. Quantification of these phenotypes was performed with the ImageJ software. **** *p*-value<0.0001, using the unpaired Student’s *t*-test (**B**). For each genotype, the percentage of embryos with each staining category was recorded. *** *p*-value<0.001, ns: non significant, using the χ2 test (**C**). (**D**) Model of Aub-dependent translational activation. See text for details.

## Discussion

Several studies have reported the role PIWI proteins in cellular mRNA regulation at the level of stability. piRNA-dependent binding of mRNAs by PIWI proteins leads to their decay in different biological systems (*16*). In addition, in *Drosophila* embryos, mRNA binding by the PIWI protein Aub also leads to their stabilization in a spatially regulated manner (*13*). Here, we report a novel function of Aub in direct translational control of mRNAs. Using *nos* mRNA as a paradigm, we show that Aub is required for *nos* mRNA translation. Nos protein levels are also strongly reduced in *armi* mutant, in which piRNA biogenesis is massively affected (*27*), suggesting that Aub loading with piRNAs is necessary for its function in translational activation. Importantly, Nos levels are not affected in a *panx* mutant background. Panx is a piRNA factor required for transcriptional repression of transposable elements, but which has no function in piRNA biogenesis (*28, 29*). In addition, as is the case for *aub* and *armi* mutants, *panx* mutant embryos do not develop (*28, 29*). Finally, Nos levels are similar in unfertilized eggs and embryos overexpressing Osk, demonstrating that Nos protein synthesis is independent of embryonic development. Together these results strongly argue for a direct role of Aub and piRNAs in *nos* mRNA translational control, independently of their role in transposable element regulation or developmental defects in piRNA pathway mutants.

Mass spectrometry analysis of Aub interactors points to a strong link with the translation machinery. In addition, with polysome gradient analyses reveal Aub association with actively translated mRNAs in polysomal fractions. A link has been reported previously between the PIWI proteins Miwi and Mili and the translation machinery in mouse testes, where Miwi and Mili were found to associate with the cap-binding complex (*46, 47*). However, the role of Miwi and Mili in translational control has not been characterized. We now decipher the molecular mechanisms of Aub function in translational activation of germ cell mRNAs in the *Drosophila* embryo. We demonstrate a physical interaction between Aub and the translation initiation factors PABP, eIF4E and subunits of the eIF3 complex. These interactions are in agreement with polysome gradient analyses in wild-type and *aub* mutant backgrounds, showing the role of Aub in translation initiation.

Recent data have identified specific roles of eIF3 in the regulation of translation. eIF3 is the most elaborate of translation initiation factors containing twelve subunits and an associated factor, eIF3j. This complex promotes all steps of translational initiation and does so in part through direct association with other translation initiation factors, contributing to their functional conformations on the small ribosomal subunit surface (*39*). In addition to this role in basal translation, the eIF3a, b, d and g subunits were shown to directly bind 5’UTR of specific mRNAs, leading to cap-dependent translation activation or repression (*40*). The eIF3d subunit that attaches to the edge of the complex appears to play an especially important role in various modes of eIF3-dependent translational control: 1) eIF3d is involved in the translational repression of *Drosophila sex-lethal* mRNA through binding to its 5’UTR (*48*). 2) eIF3d was reported to directly bind the cap structure of specific mRNAs in mammalian cells, thus bypassing the requirement of eIF4E binding to the cap for translation initiation (*41*). 3) In the same lines, eIF3d was involved in cap-dependent translational activation of specific mRNAs for neuronal remodeling in *Drosophila* larvae, in a context where eIF4E is blocked by 4E-binding protein (4E-BP) (*49*).

Other studies have reported the role of eIF3 in promoting cap-independent translation, thus highlighting eIF3 functional versatility in the control of translation. eIF3 was shown to directly bind methylated adenosines m^6^A, in mRNA 5’UTRs to induce cap-independent translation under stress conditions (*50*). Furthermore, PABP bound to the poly(A) tail was also shown to cooperate with eIF3 for its binding to mRNA 5’UTR triggering cap-independent translation (*51*).

Here, we described a new mode of eIF3-dependent translational activation through its recruitment by the PIWI protein Aub. Based on previous information on the *nos* translation repressor complex and data presented here on translational activation, we propose the following model (Fig. 6D). *nos* mRNA translation is repressed in the somatic part of the embryo by two mechanisms (*11, 35*). First, the 4E-BP protein Cup in complex with Smg binds to eIF4E and prevents eIF4G recruitment and cap-dependent translation (*11, 52*). The detailed mechanism of Cup recruitment to the repressor complex has not been clarified, but Cup was shown to directly associate with the Not1 subunit of the CCR4-NOT complex and this interaction might stabilize Cup association to eIF4E (*53*). CCR4-NOT itself is recruited to *nos* mRNA by Smg and Aub (*9, 18*). Second, two translational repressors, the RNA helicase Me31B (*Drosophila* DDX6) and its partner Tral coat the length of *nos* mRNA and prevent translation through a cap-independent mechanism (*35*). Again the mode of Me31B/Tral specific recruitment to *nos* mRNA has not been determined but the CCR4-NOT complex might also be involved since DDX6 directly binds the Not1 subunit of CCR4-NOT (*54, 55*). In the germ plasm, Osk interaction with Smg prevents Smg binding to *nos* mRNA (*9*) and this contributes to CCR4-NOT displacement from the mRNP complex. Consistent with this, CCR4 is depleted in the germ plasm (*13*). The lack of CCR4-NOT on *nos* mRNA might preclude the recruitment of Me31B/Tral and relieve the cap-independent mechanism of translational repression (Fig. 6D). We find that Aub physically interacts with PABP and several subunits of eIF3. We propose that these associations would lead to translational activation independently of eIF4E through binding of eIF3 to *nos* 5’UTR, followed by direct recruitment of the 40S ribosome by eIF3 and PABP, as previously reported for translation of *XIAP* mRNA (*51*). Alternatively, eIF3 might act through direct binding of eIF3d to the cap structure; however, we do not favor this hypothesis. Indeed, if eIF3d interaction with the cap was involved, overexpression of the point mutant eIF3d^helix11^ that is unable to bind the cap (*41*), would be expected to induce negative dominant defects, due to the lack of translation mediated by this interaction (*49*). However, overexpression of eIF3d^helix11^ with the *nos-Gal4* driver did not induce any defects in embryonic development or Nos protein synthesis (Fig. S6).

Germ granules coordinate germ cell mRNA regulation with piRNA inheritance through the role of PIWI proteins in both processes. Recent studies in *C. elegans* have shown that piRNA/PRG1-dependent mRNA accumulation in germ granules prevent their silencing, strengthening the function of piRNAs in germ granules for mRNA storage and surveillance (*56, 57*). In *Drosophila*, Aub mediates the link between piRNAs and mRNA regulation in germ granules since Aub localization to germ granules depends on its loading with piRNAs (*12*) and Aub/piRNAs play a general role in the localization and stabilization of germ cell mRNAs in germ granules (*13, 19*). How do germ granules accommodate translational control has remained more elusive. In *Drosophila* embryos, germ granules contain mRNAs that are translated sequentially (*58*). We demonstrate a direct role of Aub in translational activation. Strikingly, PABP and eIF3d tend to colocalize with Aub at the periphery of germ granules. This is reminiscent of a study analyzing translational control in relation to RNA granules in *Drosophila* oocytes, in which translational repressors such as Me31B were found to concentrate in the granule core with repressed mRNAs, whereas the translational activator Orb was localized at the edge of the granules where mRNAs docked for translation (*59*). Similarly, germ granules in embryos might be partitioned into functional subdomains involved in various steps of mRNA regulation, including storage (in an internal region of granules) and translational activation (at the granule periphery). Our work reveals the central role of Aub in activation of translation. Future studies will undoubtedly address the complexity of mRNA regulation by PIWI proteins in relation with germ granules.

## Materials and Methods

Materials and Methods are provided in the Supplementary Materials.

The mass spectrometry proteomics data have been deposited to the ProteomeXchange Consortium via the PRIDE (*60*) partner repository.

## Acknowledgements

We thank S Dorner, A Nakamura, P Lasko, M Siomi, C Smibert and A Vincent for their gifts of antibodies, and A Ephrussi, F Gebauer, S Rumpf and E Wahle for their gifts of fly stocks and DNA clones. This work was supported by the CNRS-University of Montpellier UMR9002, ANR (ANR-15-CE12-0019-01), FRM (“Equipe FRM 2013 DEQ20130326534”). AR was supported by the Labex EpiGenMed/University of Montpellier (ANR-10-LABX-12-01) and from the Fondation ARC, and CJ from the Labex EpiGenMed. MRGS, JD, CG, JC and VP held a salary from ANR, and RNS from AFM-Telethon and FRM.

## Author contribution

AR, MRGS, CJ, RNS, JD, CG, AC conducted the experiments and analyzed the data; VP and JC performed bioinformatic analyses; MD performed the mass spectrometry analysis; AB helped with polysome gradients; FJ produced clones and provided advices for LUMIER assays. MS designed the study. MS and AR wrote the manuscript; all authors commented on the manuscript.

## Materials and Methods

### Drosophila lines

*w^1118^* was used as a control. Mutant alleles and transgenic lines are *aub^QC42^ cn^1^bw^1^/CyO* and *aub^HN2^cn^1^bw^1^/CyO* (*1*), *nos-Gal4-VP16* (*2*), *UASp-osk-K10* (*3*), *panx^M1^* and *panx^M4^* (*4*), *armi^1^* (*5*), *armi^72.1^* (*6*), *w; osk^54^ nos-Gal4-VP16/TM3 Sb* and *yw; osk^54^ e UASp-GFP-Aub/TM3 Sb* (*7*), *UASp-GFP-Aub* (*8*), *UASp-GFP-Aub^AA^* (*7*), *UAS-GFP cytoplasmic* (Gift from J.M. Dura), *eIF3d^EY05735^* (Bloomington Drosophila Stock Center #20072), *eIF3d^EP-654a^* (Bloomington Drosophila Stock Center #43437). The *UASp-HA-eIF3d* and *UASp-HA-eIF3d*^helix11^ lines were generated in this study and inserted by PhiC31 recombination into attP40 site (BestGene). The genotype of embryos (aged 0-2 hours) indicated throughout are the genotypes of mothers. Females of the indicated genotypes were crosses with wild-type males.

### S2R+ cells

S2R+ cells (Gift from G. Cavalli) were cultivated at 25°C in Schneider medium complemented with 10% Fetal Bovine Serum (Gibco) and 1% penicillin-streptomycin (Gibco).

### Immunostaining and image analysis

0-2 hour-embryos were collected in a basket from plates, washed in tap water and dechorionated using commercial bleach for 2 minutes, rinsed and dried. Embryos were then fixed at the interface of a 1:1 solution of 36% formaldehyde: 100% heptane for 5 min, followed by 100% methanol devitellinisation.

Embryos were re-hydrated, blocked in 1% BSA for 1 hour and incubated overnight with primary antibodies. Secondary antibody incubation, after washes in PBS, 0.1% tween, was performed for 1 h at room temperature. Embryos were mounted in Vectashield (Vector Laboratories) for imaging. Antibodies used: rabbit anti-Osk (1/1000, Gift from P. Lasko), rabbit anti-Nos (1/1000, Gift from A. Nakamura), rabbit anti-PABP (1/500, Gift from A. Vincent), mouse anti-HA (1/2000, ascites produced from clone 12CA5), rabbit anti-GFP (1/1000, Invitrogen) and mouse anti-GFP (1/1000, Roche), goat anti-mouse IgG Cy3 (1/1000, Jackson ImmunoResearch), goat anti-rabbit IgG Alexa-488 (1/800, Invitrogen) and donkey anti-rabbit Cy3 (1/1000, Jackson ImmunoResearch). Microscopy was performed using a Leica-SP8 CLSM. Data was processed and analyzed using the ImageJ software.

### Single molecule fluorescence *in situ* hybridization (smFISH)

Dechorionated embryos were fixed at the interface of a 1:1 solution of 10% formaldehyde:100% heptane for 20 min, followed by 100% methanol devitellinisation. After permebilisation in ethanol, embryos were washed 4 times for 15 min in PBT and then once 20 min in Wash Buffer (10% 20X SCC, 10% Formamide). They were then incubated overnight at 37°C in Hybridization Buffer (10% Formamide, 10% 20X SSC, 400µg/mL tRNA, 5% Dextran sulfate, 1% VRC (Vanadyl Ribonucleoside Complexes, Sigma) and anti-*nos* probes coupled to Quasar 590 (Stellaris)). Embryos were washed in Wash Buffer at 37°C and then in 2X SCC, 0.1% Tween at room temperature before mounting (Pro-Long® Gold antifade reagent, Invitrogen). Microscopy was performed using a Leica-SP8 CLSM. Data was processed and analyzed using the ImageJ software.

### RNA extraction and RT-qPCR

Total RNA was prepared from 30 embryos using Trizol (Invitrogen) following recommendations from the manufacturer. For RT-qPCR, 1μg of total RNA was reverse transcribed using Superscript III (Invitrogen) and random hexamers. Quantitative PCR (q-PCR) was performed on a LightCycler LC480 (Roche) with Lightcycler 480 SYBR green master (Roche) and primers listed in Table S2. Quantifications were performed in triplicate.

### Coimmunoprecipitations and western blots

GFP immunoprecipitations for mass spectrometry were performed as follows. 0.5 g of 0-2 hour-dechorionated embryos were crushed in DXB buffer (25mM Hepes, 250mM Sucrose, 1mM MgCl2, 1mM DTT, 150mM NaCl, Protease inhibitor) with 0.1% Triton X-100 and RNasin and incubated on ice for 30 min. Lysates were centrifuged for 10 min and the supernatant was transferred to a new tube. Lysates were incubated on equilibrated GFP-trap beads (Chromotek), overnight at 4°C, on a wheel. Beads were washed seven times in DXB buffer complemented with 1% Triton X-100 and RNasin. Beads were suspended in 2X NuPAGE Blue supplemented with 50 mM DTT and incubated for 10 min at 95°C. The quality of the samples was assessed by silver staining (SilverQuest, Invitrogen). For co-immunoprecipitation experiments, 0.15 to 0.18 mg of 0-2 hour-embryos were crushed in IP buffer (20mM TRIS pH 7.5, 150mM NaCl, 0.2% NP-40, 1.5mM DTT, 10mM EGTA, Protease inhibitor) with either 40U/µL RNase A, or 100u/µL RNase inhibitor. Extracts were centrifuged at 10.000xg for 10 min at 4°C and incubated on pre-equilibrated magnetic beads with anti-GFP (Chromotek) or anti-HA (Pierce) for 2.5 h at 4°C. After incubation, the beads were washes five times with IP buffer and immunoprecipitated proteins were eluted from beads with 2X Laemmli buffer supplemented with 10% β-mercaptoethanol and incubated for 5 min at 95°C. Samples were then analyzed on western blots. For western blot analysis, protein extracts obtained from 30 embryos crushed in 30µl of 2X Laemmli buffer supplemented with 10% β-mercaptoethanol were boiled 5 min at 95°C. Samples were then loaded onto 10% SDS-PAGE gels before transfer to a nitrocellulose membrane. The membrane was blocked 1 hour in 5% milk diluted in PBS 1X, 0.1% Tween 20 before proceeding to primary antibody incubation (overnight, 4°C on a rotating plate). Antibody dilutions for western blots were: rabbit anti-Nos (1/1000, Gift from A.Nakamura), rabbit anti-Osk (1/2000, Gift from P. Lasko), mouse anti-Aub (4D10, 1/5000, (*9*)), guinea pig anti-Smg (1/2000, Gift from C. Smibert), rabbit anti-PABP (1/500, Gift from A. Vincent), rabbit anti-GFP (1/1000, Invitrogen), anti-HA (1/1000, Covance) and mouse anti-αTubulin (1/5000, Sigma). After washes in PBS 1X, 0.1% Tween 20, the membrane was incubated for 1 h at room temperature with secondary antibody coupled with HRP (Jackson ImmunoResearch). After washes, HRP-conjugated secondary antibodies were revealed by chemiluminescent detection (Pierce). Quantifications were performed with the ImageJ software using the Gels tool.

### Mass spectrometry

Total protein elutions were loaded on 10% SDS-PAGE gels (Mini-Protean TGX Precast gels, Bio-Rad). For each sample, one band was cut after stacking migration. Gel pieces were destained with three washes in 50% acetonitrile and 50 mM TEABC (trimethy ammonium bicarbonate buffer). After protein reduction (10 mM DTT in 50 mM TEABC at 60°C 30 min) and alkylation (55 mM iodoacetamide in TEABC at room temperature in the dak for 30 min), proteins were in-gel digested using 1µg Trypsin (Trypsin Gold, Promega). Digested products were dehydrated in a vacuum centrifuge. Obtained peptides were analyzed online using Q-Exactive Plus mass spectrometer (Thermo Fisher Scientific) interfaced with a nano-flow HPLC (RSLC U3000, Thermo Fisher Scientific). Samples were loaded onto a 15 cm reverse phase column (Acclaim Pepmap 100^®^, NanoViper, Thermo Fisher Scientific) and separated using a 103 min gradient of 2 to 40% of buffer B (80% acetonitrile, 0.1% formic acid) at a flow rate of 300 nL/min. MSMS analyzes were performed in a data-dependant mode (Xcalibur software 4.1, Thermo Fisher Scientific). Full scans (375-1500 m/z) were acquired in the Orbitrap mass analyzer with a 70,000 resolution at 200 m/z. The twelve most intense ions (charge states ≥ 2) were sequentially isolated and fragmented by HCD (high-energy collisional dissociation) in the collision cell and detected at 17,500 resolution. The spectral data were analyzed using the Maxquant software (v1.5.5.1) with default settings (*10*). All MS/MS spectra were searched by the Andromeda search engine against a decoy database consisting in a combination of *Drosophila melanogaster* entries from Reference Proteome (UP000000803, release 2018_02, https://www.uniprot.org/), isoform C sequence of Aub protein and classical contaminants, containing forward and reverse entries. Default search parameters were used, Oxidation (Met) and Acetylation (N-term) as variable modifications and Carbamidomethyl (Cys) as fixed modification were applied. FDR was set to 1% for peptides and proteins. A representative protein ID in each protein group was automatically selected using in-house bioinformatics tool (Leading_v3.2). First, proteins with the most numerous identified peptides are isolated in a “match group” (proteins from the “Protein IDs” column with the maximum number of “peptides counts”). For the match groups where more than one protein ID are present after filtering, the best annotated protein in UniProtKB, release 2019_01 (reviewed entries rather than automatic ones, highest evidence for protein existence) is defined as the “leading” protein. Label free quantification (MaxQuant LFQ) was used to identify differential proteins between samples.

### LUMIER assays

S2R+ cells (250 000) were transfected using Effectene transfection reagent (Qiagen) and incubated for 48 h at 25°C. Cells were lysed in HNTG buffer (Hepes 20mM, NaCl 150mM, MgCl2 1mM, EGTA 1mM, Triton 1%, glycerol 10%) complemented with protease inhibitor (cOmplete^TM^ EDTA-free Protease Inhibitor Cocktail, Roche). On a plate (LUMITRAC 600 96W Microplate High Binding, Greiner) pre-coated with anti-FLAG antibody (Sigma, F1804) and blocked for 1 h with a blocking solution (3% BSA, 5% sucrose and 0.5% Tween 20), lysates were incubated for 3 h on ice. After washes with HNTG, luminescence was revealed using Dual-Luciferase® Reporter Assay System (Promega, E1910) and read on a luminometer Tristar LB941. Transfections were repeated 8 to 32 times.

### Polysome profiling

For lysis of embryos we used either of two methods that produce similar results. Either 0.2 g of fresh 0-2 hour-embryos were homogenized in lysis buffer composed of 20 mM Tris-HCl, 140 mM KCl, 5 mM MgCl2, 0.5 mM DTT, 1% Triton-X100, 0.02 U/µL RNasin, 1x protease inhibitor and 0.1 mg/mL cycloheximide, or 0.2 g of frozen 0-2 hour-embryos were homogenized in lysis buffer composed of 30 mM tris-HCl, 100 mM NaCl, 10 mM MgCl2, 0.5 mM DTT, 1% Triton-X100, 0.02 U/µL RNasin, 1x protease inhibitor and 0.1 mg/mL cycloheximide. Embryo extracts were incubated 30 min on ice. Homogenates were cleared by full speed centrifugation for 30 min at 4°C. For treatment with puromycin, fresh embryos were homogenized in lysis buffer composed of 20 mM Tris-HCl, 140 mM KCl, 0.5 mM DTT, 1% Triton-X100, 0.02 U/µL RNasin, 1x protease inhibitor and 2 mM puromycin. Extracts were incubated 20 min on ice followed by 20 min at 37°C and cleared as before. The amount of RNA in the extracts was quantified using Nanodrop. Volumes of extract containing equal amounts of RNA were loaded on top of 10-50% sucrose gradient containing cycloheximide except for puromycin-treated samples. Gradients were centrifuged for 2 hours at 34,000 rpm in a SW41 rotor, with no brake. 1 mL fractions were collected using a ISCO gradient collector. 900 µL of each fraction were used for RNA extraction and 100 µL for protein precipitation. For RNA extraction, 500 pg of luciferase RNA was added to 900 µL of fraction to control RNA extraction. 0.5 % SDS and 10 mM EDTA was then added to each fraction, followed by a 5 min incubation. Then, RNAs were prepared using acid phenol-chloroform. 1 volume of acid phenol-chloroform was added, the mixture was vortexed and centrifuged at 12000 g for 15 min. The aqueous phase was put in a new tube with 2 µL of glycoblue and 1 volume of isopropanol, incubated for 15 min and centrifuged 15 min at 12,000 g. The RNA pellet was washed twice with 1 volume of ethanol 75%. The RNA pellet was dried for 5 min and resuspended in 20 µL of RNase-free H2O. 5 µL of each RNA sample were used for cDNA synthesis using superscript III and random primers. cDNAs were diluted at 1/10 for qPCR reactions that were performed using LightCycler 480 (Roche) and the primers listed in Table S2. Data were analyzed using the ΔCp method (*11*). For protein preparation, 400 µL of methanol and 100 µL of chloroform were added to 100 µL of fraction and vortexed. 300 µL of H2O were then added, vortexed and centrifuged at full speed for 5 min. The upper phase was discarded, 435 µL of methanol were added and the mixture was centrifuged at full speed for 5 min. The protein pellet was dried for 1 to 2 min and resuspended in 100 µL of 2X Laemmli buffer supplemented with 10% β-mercaptoethanol. Proteins were analyzed on western blots; antibody dilutions were anti-Aub (mouse 4D10, 1/5000, (*9*)), anti-PABP (rabbit, 1/1000, Gift from A. Vincent), anti-RpL10Ab (1/5000, (*12*)) and anti-RpS3 (1/1000, (*12*)).

### Cloning and recombineering

To produce the *UASp-HA-eIF3d* and *UASp-HA-eIF3d^helix11^* transgenes, eIF3d and eIF3d^helix11^ coding sequences were amplified by PCR from clones provided by S. Rumpf (*13*). PCR fragments were cloned into pENTR by directional Topo cloning (Invitrogen). The generated plasmids were used in gateway cloning to insert the sequences into pPHW (UASp-HA-attR1-ccdB-attR2-SV40 3’UTR) in which an attB (pPHW-attB) site has been inserted. The resulting fragments were then inserted in the *Drosophila* genome by PhiC31 recombination into the attP40 site (BestGene).

To produce FLAG-FFL-Aub, Aub coding sequence was PCR amplified from the p8161 plasmid (*8*) and cloned into pENTR by directional Topo cloning (Invitrogen). The resulting plasmid was used in gateway cloning to insert Aub sequence into pAFW in which the firefly luciferase (FFL) coding sequence has been added (pAct-FLAG-Firefly-RfA). To produce HA-Renilla (RL) tagged versions of eIF3b (DGRC, FI08008), eIF3d (*13*), eIF3f (DGRC, LD47792), eIF3k (DGRC, LD03569) and eIF4E (from E. Wahle), the coding sequences were amplified by PCR and cloned into pENTR by directional Topo cloning. The resulting plasmids were used in gateway cloning to insert the coding sequences into pAHW in which the RL coding sequence has been added (pAct-HA-Renilla-RfA). For the HA-RL tagged versions of PABP (DNASU, DmCD00772781), eIF3g (DNASU, DmCD00766429), eIF3h (DNASU, DmCD00764259) and eIF4a (DNASU, DmCD00764657), plasmids were directly used in gateway cloning to insert the sequence into pAHW in which the RL coding sequence has been added (pAct-HA-Renilla-RfA). FLAG-FFL-Cherry, HA-RL-Cherry and Sd-RL-HA were used as negative controls. To produce GST-PABP clones, the coding sequence of five *pAbp* domains, RRM1, RRM2, RRM3, RRM4 and PABC were amplified by PCR from a plasmid provided by E. Wahle. A stop codon (TAA) was added at the end of each domain. The different fragments were cloned into pGEX-4T-1 (Sigma) digested with *Eco*RI and *Xho*I. The plasmids containing HA-Aub(1–482) and HA-Aub(476–866) fragments were generated previously (*14*). The primers used to generate the constructs are listed in Table S2 and the list of constructs is in Table S3.

### GST pull-down assays

The plasmids containing GST-RRM1, GST-RRM2, GST-RRM3, GST-RRM4 and GST-PABC were introduced in *E. coli* BL21. Protein production was induced by IPTG treatment overnight at 18°C, or at 37°C for GST-RRM2. GST-fused proteins were affinity-purified on glutathione-Sepharose 4B beads (GE Healthcare); the beads were incubated overnight at 4  °C in PBT, cOmplete^TM^ EDTA-free Protease Inhibitor Cocktail (Roche) and 5% BSA. HA-Aub proteins were synthesized *in vitro* using the TnT Coupled reticulocyte lysate system (Promega), and were incubated with immobilized GST fusion proteins in 400  µl binding buffer (50  mM Hepes pH 7.5, 500  mM NaCl, 0.2  mM EDTA, 1  mM DTT, 0.5% Nonidet P-40, cOmplete^TM^ EDTA-free Protease Inhibitor Cocktail (Roche)) containing 0.2  µg  µL^−1^ RNase A. Incubations were performed for 1 h at 4  °C followed by 30 min at room temperature. Glutathione-Sepharose beads were then washed four times with binding buffer at room temperature. Recombinant proteins were dissociated from the beads by boiling 5  min in Laemmli buffer and separated on a SDS-PAGE gel. Western blots were revealed with mouse anti-HA antibody (Covance, MMS-101R) at dilution 1/1000.

### Quantification and statistical analysis

#### Statistical analysis of mass spectrometric data

Individual LFQ values per detected peptides were first quantile normalized given the experimental condition by using the ProStar (prostar-proteomics.org) (*15*) software with the default parameter set. After normalisation an imputation step was applied in cases where only one value was missing in each condition group by replacing the missing data by the mean of the observed value for this peptide in their respective experimental condition. Then, each individual experiment was combined into one data matrix. To account for batch effects, ComBat from the R package sva was used. After quality controls, differential expression analysis was done using Reproducibility-Optimized Test Statistic (ROTS) (*16*) for each different comparison. *p*-values and FDR were extracted and plotted using self-written R scripts. Significant proteins were annotated using the FlyMine database (*17*).

#### Immunofluorescence quantification

Fluorescent images were acquired using a Leica SP8-UV confocal scanning microscope. Quantification of fluorescent signal was performed using ImageJ tool Measure.

#### Colocalization quantification

Quantification of colocalization in Fig. 2 was performed using the Imaris software. For colocalization in granules (around nuclei), spots were defined with a minimal size of 0.5 µm and a PSF correction was applied to account for confocal acquisition deformation. Spot colocalization was determined within a radius of 0.25 µm around the center of the spot. For colocalization in foci (between nuclei), spots were defined with a minimal size of 0.2 µm and a PSF correction was applied to account for confocal acquisition deformation. Spots colocalization was determined within a radius of 0.25 µm around the center of the spot. Quantification of colocalization and overlapping signals in Fig. S3 was performed using ImageJ. Lines of 5 µm were drawn across the germ plasm to obtain the intensity profiles of GFP-Aub and PABP, or GFP-Aub and HA-eIF3d; background signal was subtracted. Each peak was manually categorized as colocalized, overlapping or separated with peaks from the other channel, as depicted in Fig. S3A, D.

---

**Table S2:**
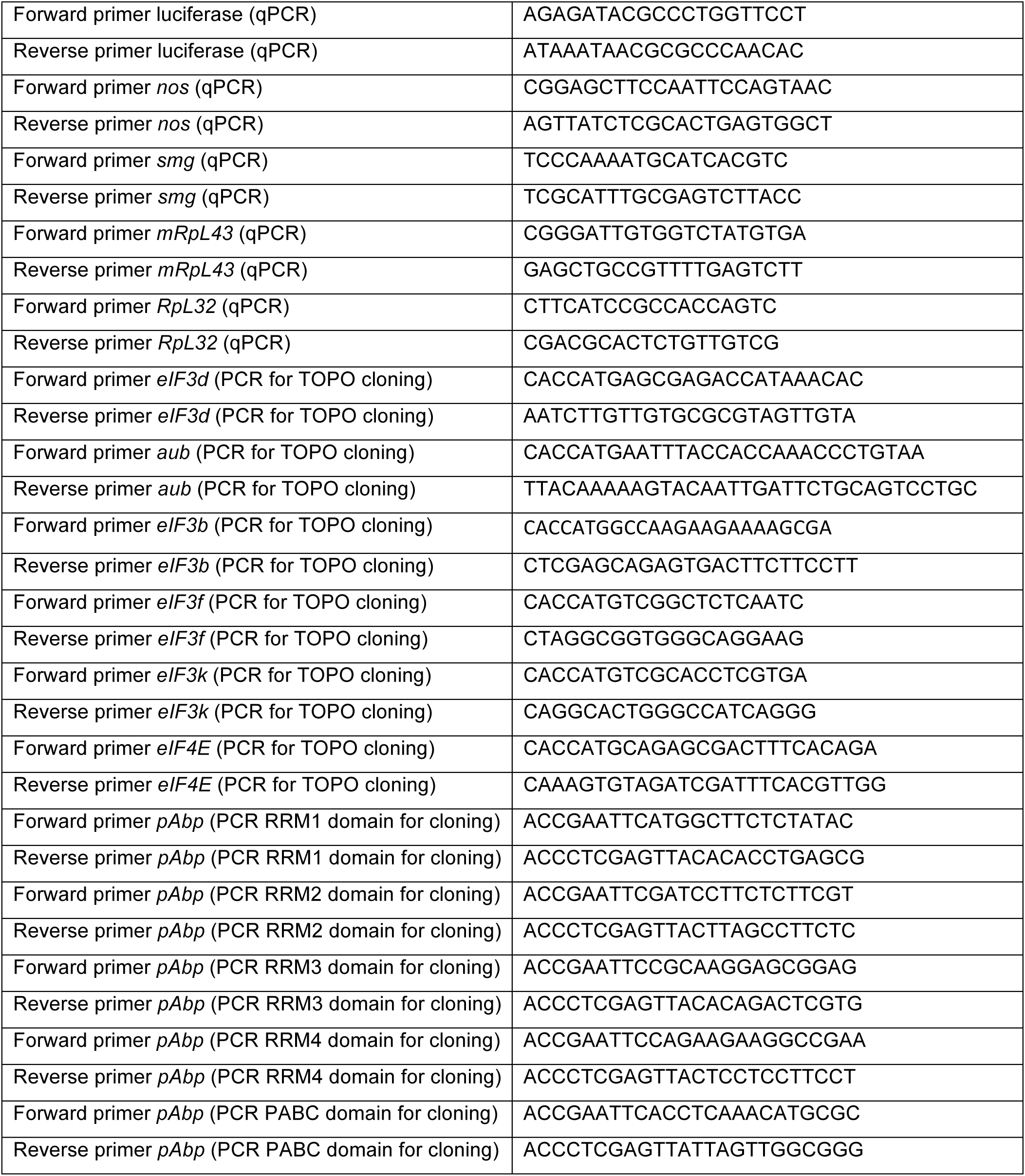
Primers used in this study.

**Table S3:** Plasmids constructed in this study.

|  |
| --- |
| pPHW-attB |
| pUASp-HA- eIF3d |
| pUASp-HA- eIF3d<sup>helix11</sup> |
| pAct-FLAG-Firefly-RfA |
| pAct-FLAG-Firefly-Cherry |
| pAct-FLAG-Firefly-Aub |
| pAct-HA-Renilla-RfA |
| pAct-Sd-Renilla-HA |
| pAct-HA-Renilla-eIF3d |
| pAct-HA-Renilla-eIF3b |
| pAct-HA-Renilla-eIF3f |
| pAct-HA-Renilla-eIF3g |
| pAct-HA-Renilla-eIF3h |
| pAct-HA-Renilla-eIF3k |
| pAct-HA-Renilla-eIF4A |
| pAct-HA-Renilla-eIF4E |
| pAct-HA-Renilla-PABP |
| pAct-HA-Renilla-Cherry |

## Supplementary Figure Legends

**Figure S1.**
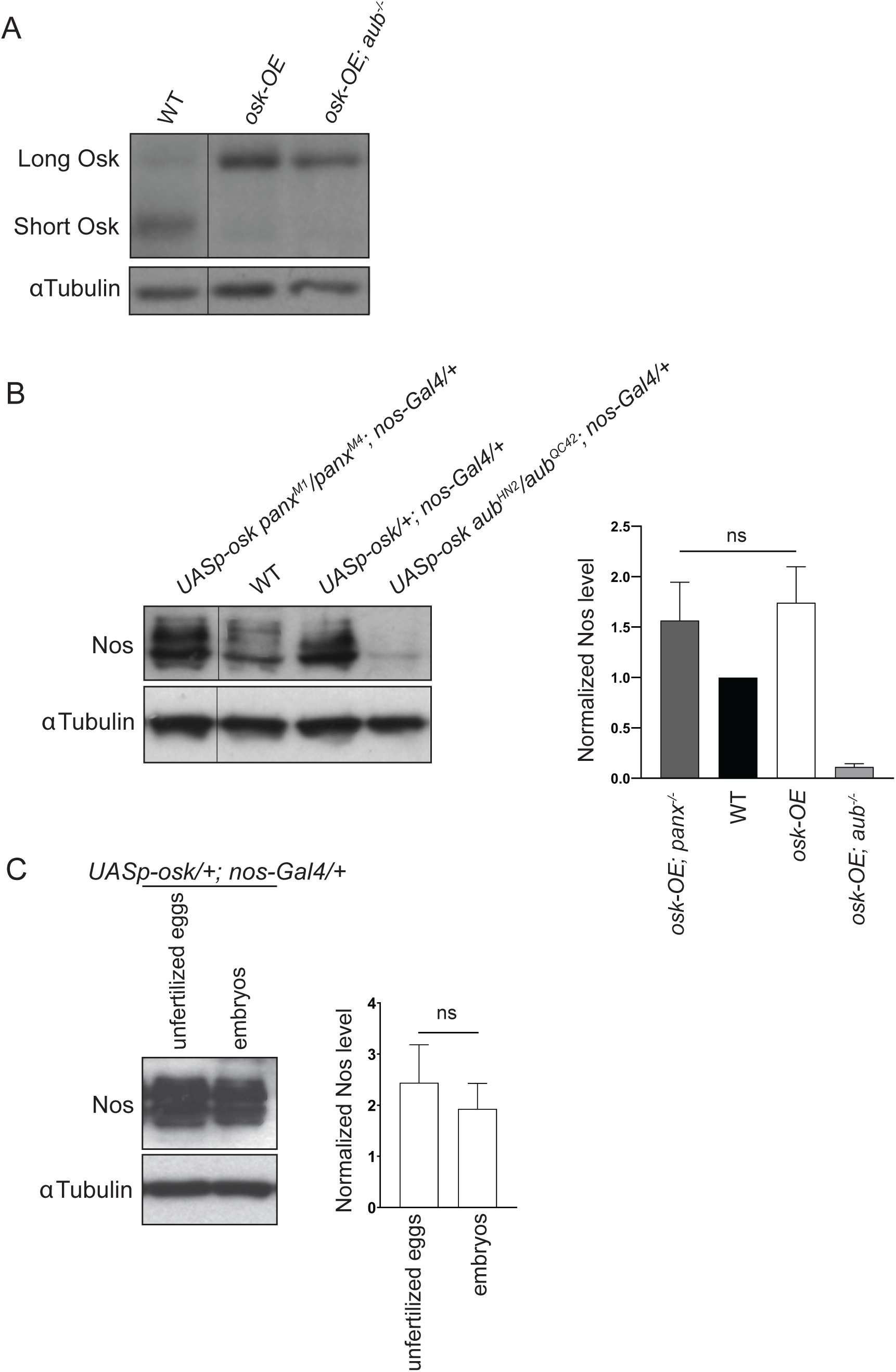
*nos* mRNA translation is independent of Panx and development. (**A**) Western blots of wild-type (WT), *osk-OE* and *osk-OE; aub^-/-^* embryos revealed with anti-Osk showing that Long Osk is overexpressed from the *UASp-osk* transgene. αTubulin was used as a loading control. (**B**) Western blots of wild-type (WT) embryos and embryos overexpressing *osk* either in a wild-type, *aub* mutant or *panx* mutant background revealed with anti-Nos, showing that Nos levels were not affected in the *panx* mutant background. αTubulin was used as a loading control. Quantification was performed using the ImageJ software with 4 biological replicates. Error bars represent sem. ns: non significant, using the unpaired Student’s *t*-test. (**C**) Western blots of *osk-OE* embryos and unfertilized eggs, showing that Nos levels did not depend on embryonic development. αTubulin was used as a loading control. Quantification was performed using the ImageJ software with 4 biological replicates. Error bars represent sem. ns: non significant, using the unpaired Student’s *t*-test.

**Figure S2.**
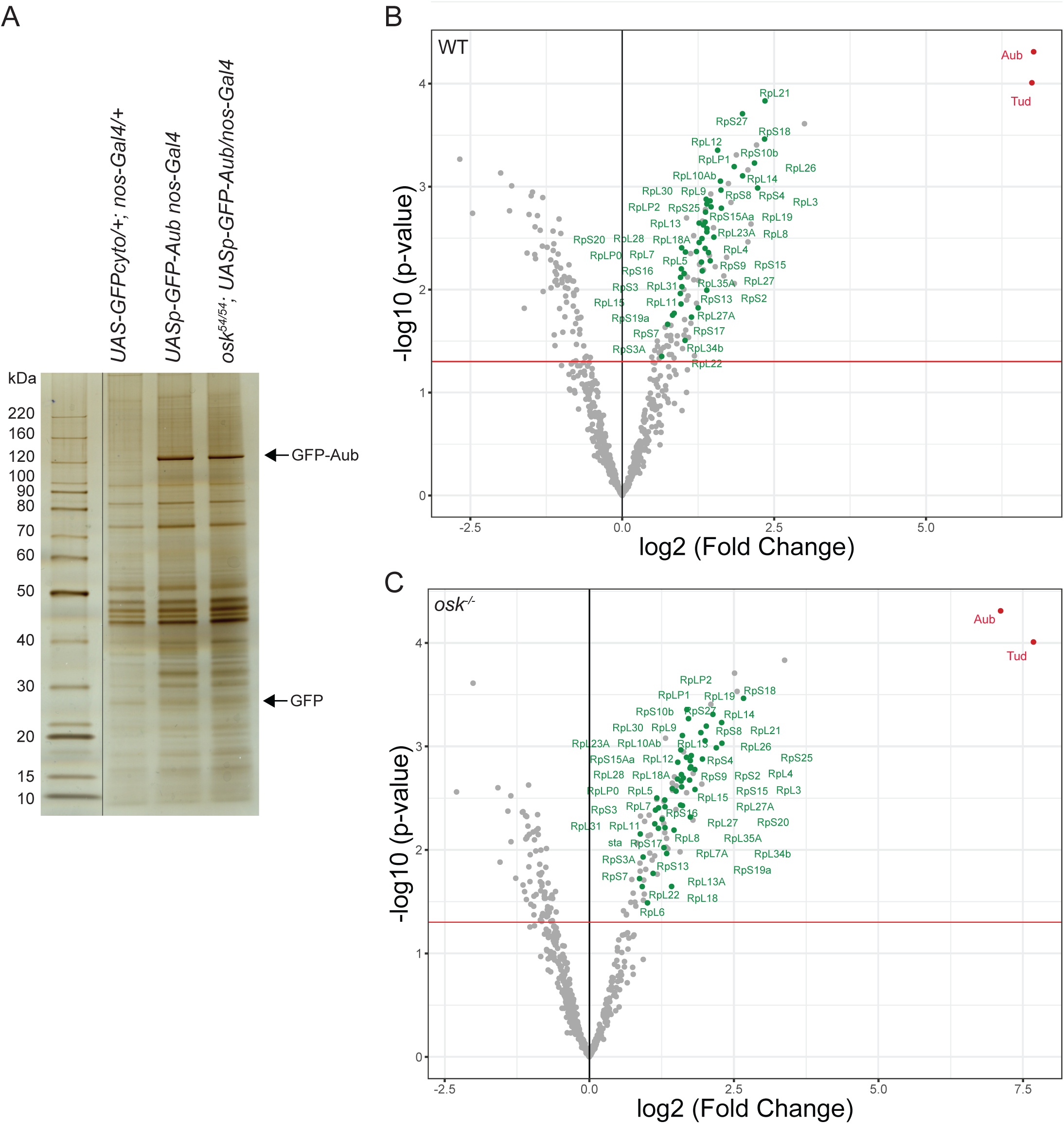
Ribosomal proteins are associated with Aub. (**A**) Silver stained gel showing protein extracts used for mass spectrometry following GFP IP. (**B, C**) Volcano plots showing the mass spectrometry analysis of GFP-Aub IP from 0-2 hour-embryos. Embryos expressing cytoplasmic GFP were used as control. (**B**) *UASp-GFP-Aub nos-Gal4* embryos; (**C**) *osk^54^; UASp-GFP-Aub/nos-Gal4* embryos. The analysis was based on four biological replicates. The red line indicates the significance threshold (*p*-value = 0.05). Ribosomal proteins are indicated in green.

**Figure S3.**
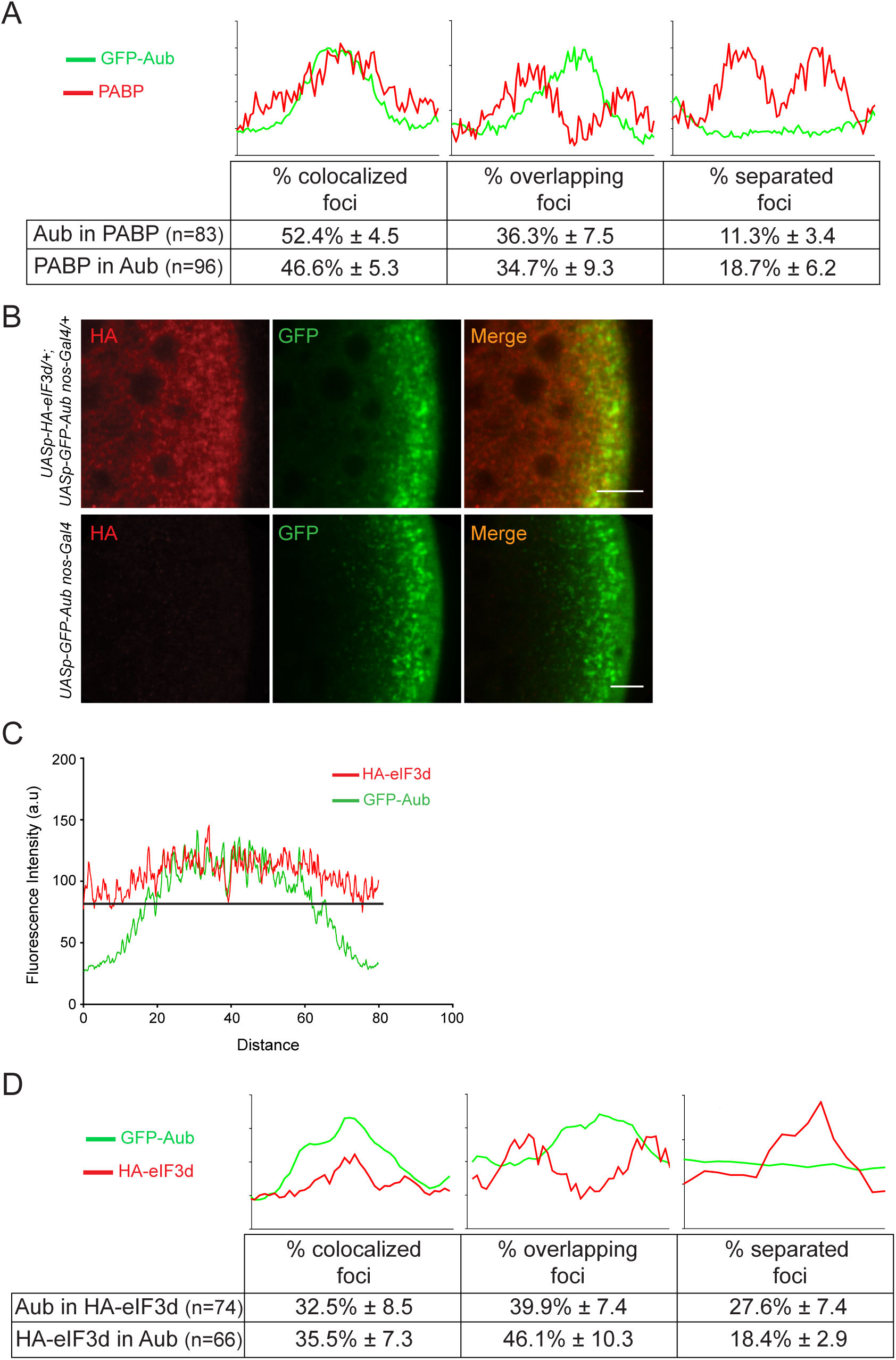
PABP and eIF3d foci colocalize or overlap with germ granules. (**A**) Quantification of colocalization and overlap between PABP foci and Aub-containing germ granules. Representative graphs for each category (colocalization; overlap; separated foci) are shown. Quantification was performed with the ImageJ software. ( **B**) Immunostaining of embryos with anti-GFP (green) to visualize Aub and anti-HA (red), showing the specificity of anti-HA antibody that did not recognize any protein in embryos lacking the *UASp-HA-eIF3d* transgene. Posterior of embryos are shown. Scale bars: 10 μm. (**C**) Graph showing the slight accumulation of eIF3d at the posterior pole of embryos. A line was drawn along the posterior cortex of the embryo shown in Figure 4I and the fluorescence intensity was quantified using the ImageJ software. (**D**) Quantification of colocalization and overlap between HA-eIF3d foci and Aub-containing germ granules. Representative graphs for each category (colocalization; overlap; separated foci) are shown. Quantification was performed with the ImageJ software.

**Figure S4.**
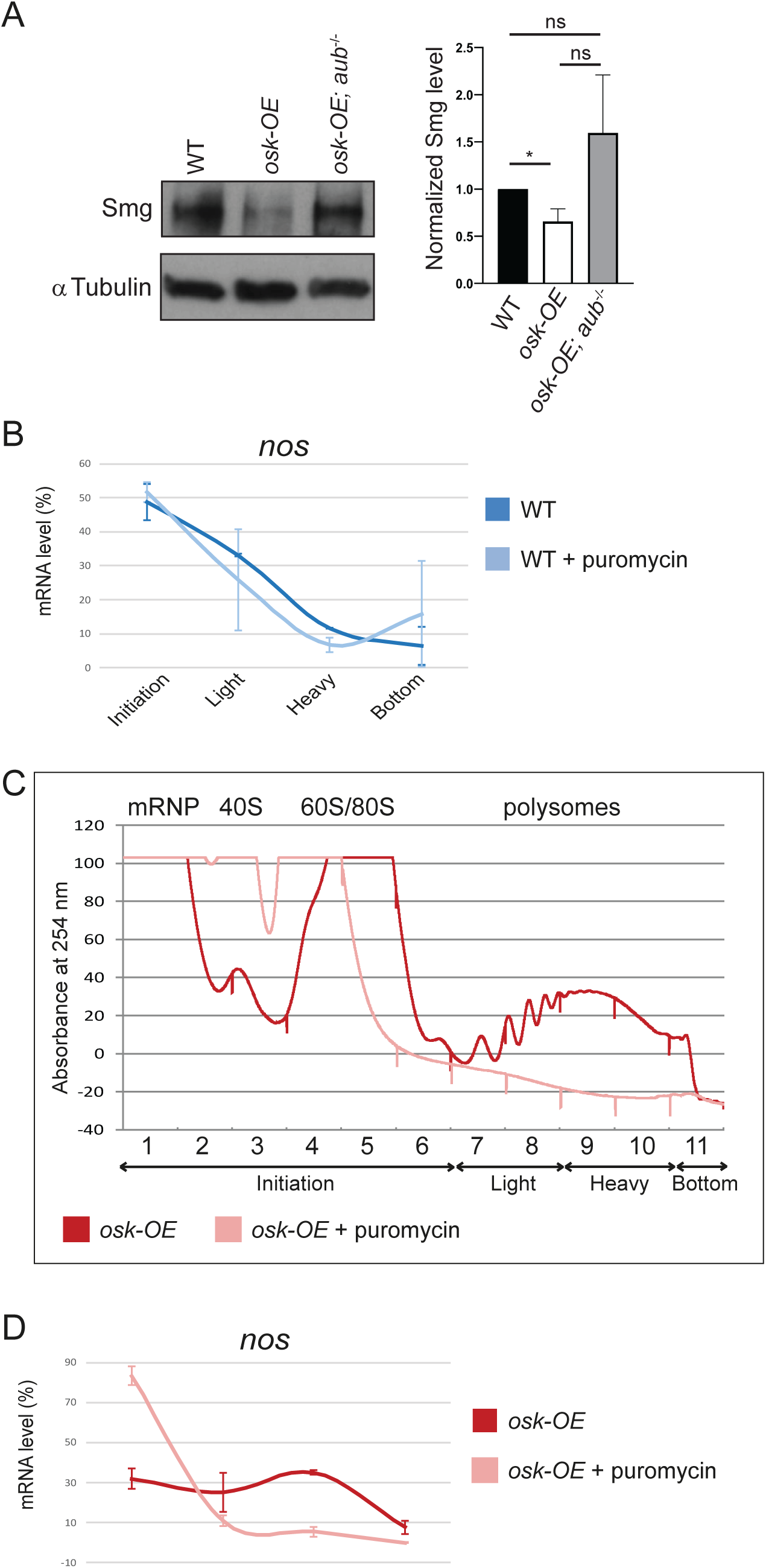
*nos* mRNA profiling through polysome gradients. (**A**) Western blots of wild-type (WT), *osk-OE* and *osk-OE; aub^-/-^* embryos revealed with anti-Smg showing that Smg levels did not decrease in *osk-OE; aub^-/-^* embryos. αTubulin was used as a loading control. Quantification was performed using the ImageJ software with 4 biological replicates. Error bars represent sem. \**p*-value<0.05, ns: non significant, using the unpaired Student’s *t*-test. (**B**) Quantification of *nos* mRNA using RT-qPCR in the different fractions of the gradients for wild-type embryos (WT) in the absence (dark blue) or the presence of puromycin (light blue). mRNA levels are indicated in percentage of total mRNA in all the fractions. Mean of two biological replicates, quantified in triplicates. Error bars represent sem. (**C**) Profile of 254 nm absorbance for 0-2 hour *osk-OE* embryos treated (pink), or not (red) with puromycin, fractionated into 10-50 % sucrose gradients. (**D**) Quantification of *nos* mRNA using RT-qPCR in the different fractions of the gradients for *osk-OE* embryos in the absence (red) or the presence of puromycin (pink). mRNA levels are indicated in percentage of total mRNA in all the fractions. Mean of two biological replicates, quantified in triplicates. Error bars represent sem.

**Figure S5.**
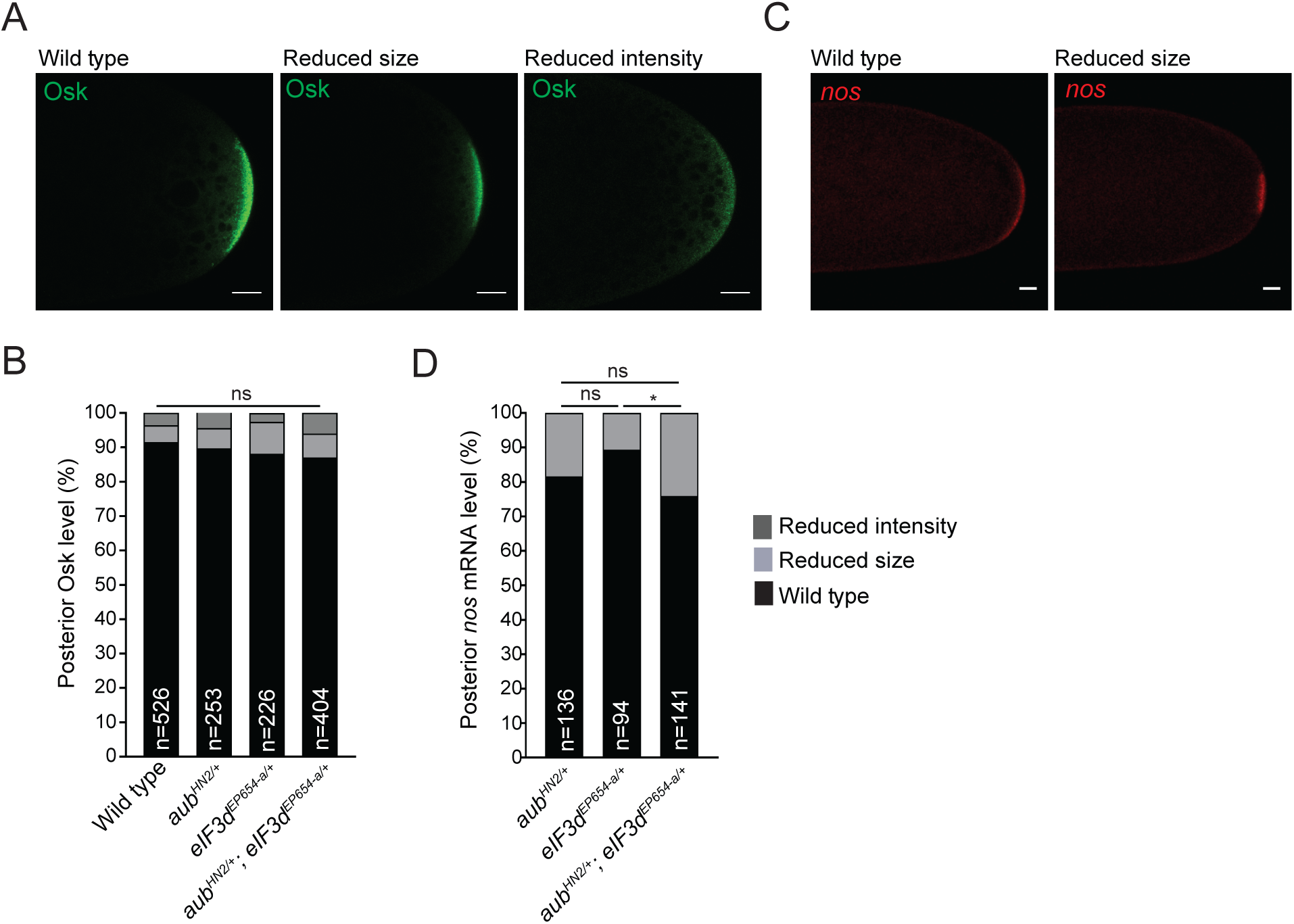
Aub-eIF3d interaction does not regulate Osk posterior accumulation. (**A, B**) Immunostaining of single and double *aub* and *eIF3d* heterozygous mutant embryos with anti-Osk antibody. Posterior of embryos showing the three types of staining: wild type, reduced size or reduced intensity (**A**). For each genotype, the percentage of embryos with each staining category was recorded. ns: non significant, using the χ2 test (**B**). (**C, D**) *nos* smFISH of single and double *aub* and *eIF3d* heterozygous mutant embryos. Posterior of embryos showing the two types of obtained staining: wild type and reduced size (**C**). For each genotype, the percentage of embryos with each staining category was recorded. * *p*-value<0.05, ns: non significant, using the χ2 test (**D**).

**Figure S6.**
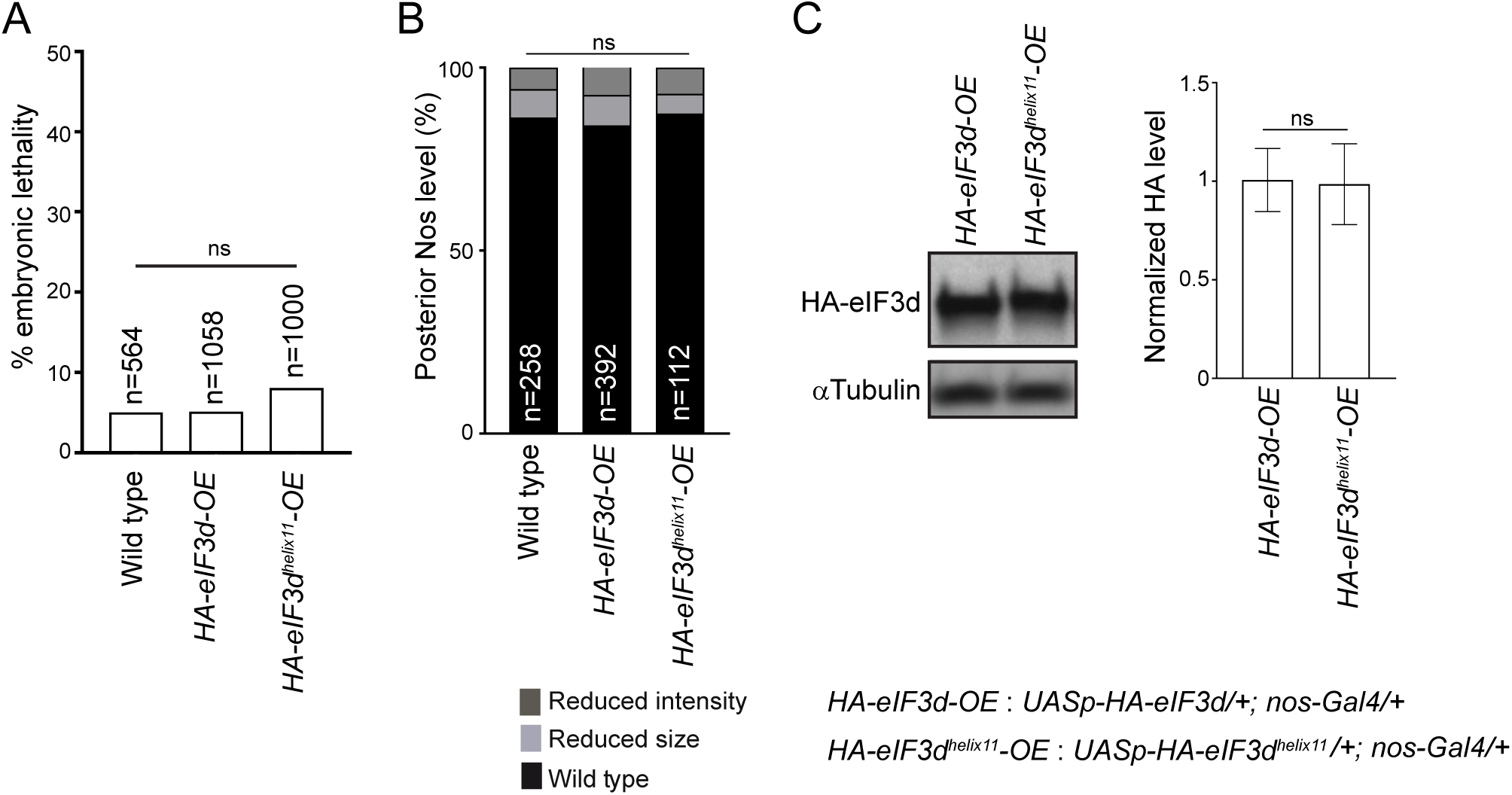
*eIF3d^helix11^* does not act as a negative dominant mutant in the embryo. (**A**) Percentage of lethality of embryos overexpressing HA-eIF3d or HA-eIF3d^helix11^. The genotypes are indicated. ns: non significant, using the χ2 test. (**B**) Quantification of immunostaining with anti-Nos antibody of embryos overexpressing HA-eIF3d or HA-eIF3d^helix11^. The three types of staining, wild type, reduced size or reduced intensity were as in Figure 6B. For each genotype, the percentage of embryos with each staining category was recorded. ns: non significant, using the χ2 test. (**C**) Western blot of embryos overexpressing HA-eIF3d or HA-eIF3d^helix11^ revealed with anti-HA, showing that the level of overexpression is similar for eIF3d and eIF3d^helix11^. αTubulin was used as a loading control. Quantification was performed using the ImageJ software with 3 biological replicates. Error bars represent sem. ns: non significant, using the unpaired Student’s *t*-test.

## References

1. B. Barckmann, M. Simonelig, Control of maternal mRNA stability in germ cells and early embryos. Biochimica et biophysica acta 1829, 714–724 (2013)

2. E. R. Gavis, R. Lehmann, Localization of nanos RNA controls embryonic polarity. Cell 71, 301–313 (1992)

3. S. E. Bergsten, E. R. Gavis, Role for mRNA localization in translational activation but not spatial restriction of nanos RNA. Development 126, 659–669 (1999)

4. T. Trcek, M. Grosch, A. York, H. Shroff, T. Lionnet, R. Lehmann, Drosophila germ granules are structured and contain homotypic mRNA clusters. Nature communications 6, 7962 (2015)

5. A. Dahanukar, R. P. Wharton, The Nanos gradient in Drosophila embryos is generated by translational regulation. Genes Dev 10, 2610–2620 (1996)

6. E. R. Gavis, R. Lehmann, Translational regulation of nanos by RNA localization. Nature 369, 315–318 (1994)

7. A. Dahanukar, J. A. Walker, R. P. Wharton, Smaug, a novel RNA-binding protein that operates a translational switch in Drosophila. Mol Cell 4, 209–218 (1999)

8. C. A. Smibert, J. E. Wilson, K. Kerr, P. M. Macdonald, smaug protein represses translation of unlocalized nanos mRNA in the Drosophila embryo. Genes Dev 10, 2600–2609 (1996)

9. S. Zaessinger, I. Busseau, M. Simonelig, Oskar allows nanos mRNA translation in Drosophila embryos by preventing its deadenylation by Smaug/CCR4. Development 133, 4573–4583 (2006)

10. A. Ephrussi, R. Lehmann, Induction of germ cell formation by oskar. Nature 358, 387–392 (1992)

11. M. Jeske, B. Moritz, A. Anders, E. Wahle, Smaug assembles an ATP-dependent stable complex repressing nanos mRNA translation at multiple levels. EMBO J 30, 90–103 (2011)

12. B. Barckmann, S. Pierson, J. Dufourt, C. Papin, C. Armenise, F. Port, T. Grentzinger, S. Chambeyron, G. Baronian, J. P. Desvignes, T. Curk, M. Simonelig, Aubergine iCLIP Reveals piRNA-Dependent Decay of mRNAs Involved in Germ Cell Development in the Early Embryo. Cell Rep 12, 1205–1216 (2015)

13. J. Dufourt, G. Bontonou, A. Chartier, C. Jahan, A. C. Meunier, S. Pierson, P. F. Harrison, C. Papin, T. H. Beilharz, M. Simonelig, piRNAs and Aubergine cooperate with Wispy poly(A) polymerase to stabilize mRNAs in the germ plasm. Nature communications 8, 1305 (2017)

14. B. Czech, M. Munafo, F. Ciabrelli, E. L. Eastwood, M. H. Fabry, E. Kneuss, G. J. Hannon, piRNA-Guided Genome Defense: From Biogenesis to Silencing. Annu Rev Genet 52, 131–157 (2018)

15. X. Huang, K. Fejes Toth, A. A. Aravin, piRNA Biogenesis in Drosophila melanogaster. Trends Genet 33, 882–894 (2017)

16. P. Rojas-Rios, M. Simonelig, piRNAs and PIWI proteins: regulators of gene expression in development and stem cells. Development 145, (2018)

17. S. R. Mani, H. Megosh, H. Lin, PIWI proteins are essential for early Drosophila embryogenesis. Dev Biol 385, 340–349 (2014)

18. C. Rouget, C. Papin, A. Boureux, A. C. Meunier, B. Franco, N. Robine, E. C. Lai, A. Pelisson, M. Simonelig, Maternal mRNA deadenylation and decay by the piRNA pathway in the early Drosophila embryo. Nature 467, 1128–1132 (2010)

19. A. Vourekas, P. Alexiou, N. Vrettos, M. Maragkakis, Z. Mourelatos, Sequence-dependent but not sequence-specific piRNA adhesion traps mRNAs to the germ plasm. Nature 531, 390–394 (2016)

20. W. S. Goh, I. Falciatori, O. H. Tam, R. Burgess, O. Meikar, N. Kotaja, M. Hammell, G. J. Hannon, piRNA-directed cleavage of meiotic transcripts regulates spermatogenesis. Genes Dev 29, 1032–1044 (2015)

21. L. T. Gou, P. Dai, J. H. Yang, Y. Xue, Y. P. Hu, Y. Zhou, J. Y. Kang, X. Wang, H. Li, M. M. Hua, S. Zhao, S. D. Hu, L. G. Wu, H. J. Shi, Y. Li, X. D. Fu, L. H. Qu, E. D. Wang, M. F. Liu, Pachytene piRNAs instruct massive mRNA elimination during late spermiogenesis. Cell Res 24, 680–700 (2014)

22. T. Kiuchi, H. Koga, M. Kawamoto, K. Shoji, H. Sakai, Y. Arai, G. Ishihara, S. Kawaoka, S. Sugano, T. Shimada, Y. Suzuki, M. G. Suzuki, S. Katsuma, A single female-specific piRNA is the primary determiner of sex in the silkworm. Nature, (2014)

23. T. Watanabe, E. C. Cheng, M. Zhong, H. Lin, Retrotransposons and pseudogenes regulate mRNAs and lncRNAs via the piRNA pathway in the germline. Genome research 25, 368–380 (2015)

24. P. Zhang, J. Y. Kang, L. T. Gou, J. Wang, Y. Xue, G. Skogerboe, P. Dai, D. W. Huang, R. Chen, X. D. Fu, M. F. Liu, S. He, MIWI and piRNA-mediated cleavage of messenger RNAs in mouse testes. Cell Res 25, 193–207 (2015)

25. V. Riechmann, G. J. Gutierrez, P. Filardo, A. R. Nebreda, A. Ephrussi, Par-1 regulates stability of the posterior determinant Oskar by phosphorylation. Nat Cell Biol 4, 337–342 (2002)

26. F. H. Markussen, A. M. Michon, W. Breitwieser, A. Ephrussi, Translational control of oskar generates short OSK, the isoform that induces pole plasma assembly. Development 121, 3723–3732 (1995)

27. C. D. Malone, J. Brennecke, M. Dus, A. Stark, W. R. McCombie, R. Sachidanandam, G. J. Hannon, Specialized piRNA pathways act in germline and somatic tissues of the Drosophila ovary. Cell 137, 522–535 (2009)

28. G. Sienski, J. Batki, K. A. Senti, D. Donertas, L. Tirian, K. Meixner, J. Brennecke, Silencio/CG9754 connects the Piwi-piRNA complex to the cellular heterochromatin machinery. Genes Dev 29, 2258–2271 (2015)

29. Y. Yu, J. Gu, Y. Jin, Y. Luo, J. B. Preall, J. Ma, B. Czech, G. J. Hannon, Panoramix enforces piRNA-dependent cotranscriptional silencing. Science 350, 339–342 (2015)

30. J. S. Khurana, J. Xu, Z. Weng, W. E. Theurkauf, Distinct functions for the Drosophila piRNA pathway in genome maintenance and telomere protection. PLoS Genet 6, e1001246 (2010)

31. S. C. Little, K. S. Sinsimer, J. J. Lee, E. F. Wieschaus, E. R. Gavis, Independent and coordinate trafficking of single Drosophila germ plasm mRNAs. Nat Cell Biol 17, 558–568 (2015)

32. T. Thomson, N. Liu, A. Arkov, R. Lehmann, P. Lasko, Isolation of new polar granule components in Drosophila reveals P body and ER associated proteins. Mech Dev 125, 865–873 (2008)

33. Y. Kirino, A. Vourekas, N. Sayed, F. de Lima Alves, T. Thomson, P. Lasko, J. Rappsilber, T. A. Jongens, Z. Mourelatos, Arginine methylation of Aubergine mediates Tudor binding and germ plasm localization. RNA 16, 70–78 (2010)

34. K. M. Nishida, T. N. Okada, T. Kawamura, T. Mituyama, Y. Kawamura, S. Inagaki, H. Huang, D. Chen, T. Kodama, H. Siomi, M. C. Siomi, Functional involvement of Tudor and dPRMT5 in the piRNA processing pathway in Drosophila germlines. EMBO J 28, 3820–3831 (2009)

35. M. Gotze, J. Dufourt, C. Ihling, C. Rammelt, S. Pierson, N. Sambrani, C. Temme, A. Sinz, M. Simonelig, E. Wahle, Translational repression of the Drosophila nanos mRNA involves the RNA helicase Belle and RNA coating by Me31B and Trailer hitch. RNA 23, 1552–1568 (2017)

36. Y. Kirino, N. Kim, M. de Planell-Saguer, E. Khandros, S. Chiorean, P. S. Klein, I. Rigoutsos, T. A. Jongens, Z. Mourelatos, Arginine methylation of Piwi proteins catalysed by dPRMT5 is required for Ago3 and Aub stability. Nat Cell Biol, (2009)

37. P. Trepte, A. Buntru, K. Klockmeier, L. Willmore, A. Arumughan, C. Secker, M. Zenkner, L. Brusendorf, K. Rau, A. Redel, E. E. Wanker, DULIP: A Dual Luminescence-Based Co-Immunoprecipitation Assay for Interactome Mapping in Mammalian Cells. Journal of molecular biology 427, 3375–3388 (2015)

38. M. Brook, J. W. Smith, N. K. Gray, The DAZL and PABP families: RNA-binding proteins with interrelated roles in translational control in oocytes. Reproduction 137, 595–617 (2009)

39. L. S. Valasek, J. Zeman, S. Wagner, P. Beznoskova, Z. Pavlikova, M. P. Mohammad, V. Hronova, A. Herrmannova, Y. Hashem, S. Gunisova, Embraced by eIF3: structural and functional insights into the roles of eIF3 across the translation cycle. Nucleic Acids Res 45, 10948–10968 (2017)

40. A. S. Lee, P. J. Kranzusch, J. H. Cate, eIF3 targets cell-proliferation messenger RNAs for translational activation or repression. Nature 522, 111–114 (2015)

41. A. S. Lee, P. J. Kranzusch, J. A. Doudna, J. H. Cate, eIF3d is an mRNA cap-binding protein that is required for specialized translation initiation. Nature 536, 96–99 (2016)

42. T. H. Beilharz, T. Preiss, Translational profiling: the genome-wide measure of the nascent proteome. Briefings in functional genomics & proteomics 3, 103–111 (2004)

43. B. Benoit, C. H. He, F. Zhang, S. M. Votruba, W. Tadros, J. T. Westwood, C. A. Smibert, H. D. Lipshitz, W. E. Theurkauf, An essential role for the RNA-binding protein Smaug during the Drosophila maternal-to-zygotic transition. Development 136, 923–932 (2009)

44. W. Tadros, A. L. Goldman, T. Babak, F. Menzies, L. Vardy, T. Orr-Weaver, T. R. Hughes, J. T. Westwood, C. A. Smibert, H. D. Lipshitz, SMAUG is a major regulator of maternal mRNA destabilization in Drosophila and its translation is activated by the PAN GU kinase. Dev Cell 12, 143–155 (2007)

45. I. Kronja, B. Yuan, S. W. Eichhorn, K. Dzeyk, J. Krijgsveld, D. P. Bartel, T. L. Orr-Weaver, Widespread changes in the posttranscriptional landscape at the Drosophila oocyte-to-embryo transition. Cell Rep 7, 1495–1508 (2014)

46. S. T. Grivna, B. Pyhtila, H. Lin, MIWI associates with translational machinery and PIWI-interacting RNAs (piRNAs) in regulating spermatogenesis. Proc Natl Acad Sci U S A 103, 13415–13420 (2006)

47. Y. Unhavaithaya, Y. Hao, E. Beyret, H. Yin, S. Kuramochi-Miyagawa, T. Nakano, H. Lin, MILI, a PIWI-interacting RNA-binding protein, is required for germ line stem cell self-renewal and appears to positively regulate translation. J Biol Chem 284, 6507–6519 (2009)

48. E. Szostak, M. Garcia-Beyaert, T. Guitart, A. Graindorge, O. Coll, F. Gebauer, Hrp48 and eIF3d contribute to msl-2 mRNA translational repression. Nucleic Acids Res 46, 4099–4113 (2018)

49. S. Rode, H. Ohm, L. Anhauser, M. Wagner, M. Rosing, X. Deng, O. Sin, S. A. Leidel, E. Storkebaum, A. Rentmeister, S. Zhu, S. Rumpf, Differential Requirement for Translation Initiation Factor Pathways during Ecdysone-Dependent Neuronal Remodeling in Drosophila. Cell Rep 24, 2287–2299 e2284 (2018)

50. K. D. Meyer, D. P. Patil, J. Zhou, A. Zinoviev, M. A. Skabkin, O. Elemento, T. V. Pestova, S. B. Qian, S. R. Jaffrey, 5’ UTR m(6)A Promotes Cap-Independent Translation. Cell 163, 999–1010 (2015)

51. N. Thakor, M. D. Smith, L. Roberts, M. D. Faye, H. Patel, H. J. Wieden, J. H. D. Cate, M. Holcik, Cellular mRNA recruits the ribosome via eIF3-PABP bridge to initiate internal translation. RNA Biol 14, 553–567 (2017)

52. M. R. Nelson, A. M. Leidal, C. A. Smibert, Drosophila Cup is an eIF4E-binding protein that functions in Smaug-mediated translational repression. Embo J 23, 150–159 (2004)

53. C. Igreja, E. Izaurralde, CUP promotes deadenylation and inhibits decapping of mRNA targets. Genes Dev 25, 1955–1967 (2011)

54. Y. Chen, A. Boland, D. Kuzuoglu-Ozturk, P. Bawankar, B. Loh, C. T. Chang, O. Weichenrieder, E. Izaurralde, A DDX6-CNOT1 complex and W-binding pockets in CNOT9 reveal direct links between miRNA target recognition and silencing. Mol Cell 54, 737–750 (2014)

55. H. Mathys, J. Basquin, S. Ozgur, M. Czarnocki-Cieciura, F. Bonneau, A. Aartse, A. Dziembowski, M. Nowotny, E. Conti, W. Filipowicz, Structural and biochemical insights to the role of the CCR4-NOT complex and DDX6 ATPase in microRNA repression. Mol Cell 54, 751–765 (2014)

56. A. E. Dodson, S. Kennedy, Germ Granules Coordinate RNA-Based Epigenetic Inheritance Pathways. Dev Cell 50, 704–715 e704 (2019)

57. J. P. T. Ouyang, A. Folkmann, L. Bernard, C. Y. Lee, U. Seroussi, A. G. Charlesworth, J. M. Claycomb, G. Seydoux, P Granules Protect RNA Interference Genes from Silencing by piRNAs. Dev Cell 50, 716–728 e716 (2019)

58. P. Rangan, M. DeGennaro, K. Jaime-Bustamante, R. X. Coux, R. G. Martinho, R. Lehmann, Temporal and spatial control of germ-plasm RNAs. Curr Biol 19, 72–77 (2009)

59. T. T. Weil, R. M. Parton, B. Herpers, J. Soetaert, T. Veenendaal, D. Xanthakis, I. M. Dobbie, J. M. Halstead, R. Hayashi, C. Rabouille, I. Davis, Drosophila patterning is established by differential association of mRNAs with P bodies. Nat Cell Biol 14, 1305–1313 (2012)

60. Y. Perez-Riverol, A. Csordas, J. Bai, M. Bernal-Llinares, S. Hewapathirana, D. J. Kundu, A. Inuganti, J. Griss, G. Mayer, M. Eisenacher, E. Perez, J. Uszkoreit, J. Pfeuffer, T. Sachsenberg, S. Yilmaz, S. Tiwary, J. Cox, E. Audain, M. Walzer, A. F. Jarnuczak, T. Ternent, A. Brazma, J. A. Vizcaino, The PRIDE database and related tools and resources in 2019: improving support for quantification data. Nucleic Acids Res 47, D442–D450 (2019)

## References

1. T. Schupbach, E. Wieschaus, Female sterile mutations on the second chromosome of Drosophila melanogaster. II. Mutations blocking oogenesis or altering egg morphology. Genetics 129, 1119–1136 (1991)

2. P. Rorth, Gal4 in the *Drosophila* female germline. Mech. Dev. 78, 113–118 (1998)

3. V. Riechmann, G. J. Gutierrez, P. Filardo, A. R. Nebreda, A. Ephrussi, Par-1 regulates stability of the posterior determinant Oskar by phosphorylation. Nat Cell Biol 4, 337–342 (2002)

4. Y. Yu, J. Gu, Y. Jin, Y. Luo, J. B. Preall, J. Ma, B. Czech, G. J. Hannon, Panoramix enforces piRNA-dependent cotranscriptional silencing. Science 350, 339–342 (2015)

5. Y. Tomari, T. Du, B. Haley, D. S. Schwarz, R. Bennett, H. A. Cook, B. S. Koppetsch, W. E. Theurkauf, P. D. Zamore, RISC assembly defects in the Drosophila RNAi mutant armitage. Cell 116, 831–841 (2004)

6. H. A. Cook, B. S. Koppetsch, J. Wu, W. E. Theurkauf, The Drosophila SDE3 homolog armitage is required for oskar mRNA silencing and embryonic axis specification. Cell 116, 817–829 (2004)

7. B. Barckmann, S. Pierson, J. Dufourt, C. Papin, C. Armenise, F. Port, T. Grentzinger, S. Chambeyron, G. Baronian, J. P. Desvignes, T. Curk, M. Simonelig, Aubergine iCLIP Reveals piRNA-Dependent Decay of mRNAs Involved in Germ Cell Development in the Early Embryo. Cell Rep 12, 1205–1216 (2015)

8. A. N. Harris, P. M. Macdonald, Aubergine encodes a Drosophila polar granule component required for pole cell formation and related to eIF2C. Development 128, 2823–2832 (2001)

9. L. S. Gunawardane, K. Saito, K. M. Nishida, K. Miyoshi, Y. Kawamura, T. Nagami, H. Siomi, M. C. Siomi, A slicer-mediated mechanism for repeat-associated siRNA 5’ end formation in Drosophila. Science 315, 1587–1590 (2007)

10. J. Cox, M. Mann, MaxQuant enables high peptide identification rates, individualized p.p.b.-range mass accuracies and proteome-wide protein quantification. Nat Biotechnol 26, 1367–1372 (2008)

11. A. C. Panda, J. L. Martindale, M. Gorospe, Polysome Fractionation to Analyze mRNA Distribution Profiles. Bio-protocol 7, (2017)

12. S. Antic, M. T. Wolfinger, A. Skucha, S. Hosiner, S. Dorner, General and MicroRNA-Mediated mRNA Degradation Occurs on Ribosome Complexes in Drosophila Cells. Mol Cell Biol 35, 2309–2320 (2015)

13. S. Rode, H. Ohm, L. Anhauser, M. Wagner, M. Rosing, X. Deng, O. Sin, S. A. Leidel, E. Storkebaum, A. Rentmeister, S. Zhu, S. Rumpf, Differential Requirement for Translation Initiation Factor Pathways during Ecdysone-Dependent Neuronal Remodeling in Drosophila. Cell Rep 24, 2287–2299 e2284 (2018)

14. J. Dufourt, G. Bontonou, A. Chartier, C. Jahan, A. C. Meunier, S. Pierson, P. F. Harrison, C. Papin, T. H. Beilharz, M. Simonelig, piRNAs and Aubergine cooperate with Wispy poly(A) polymerase to stabilize mRNAs in the germ plasm. Nature communications 8, 1305 (2017)

15. S. Wieczorek, F. Combes, C. Lazar, Q. Giai Gianetto, L. Gatto, A. Dorffer, A. M. Hesse, Y. Coute, M. Ferro, C. Bruley, T. Burger, DAPAR & ProStaR: software to perform statistical analyses in quantitative discovery proteomics. Bioinformatics 33, 135–136 (2017)

16. T. Suomi, F. Seyednasrollah, M. K. Jaakkola, T. Faux, L. L. Elo, ROTS: An R package for reproducibility-optimized statistical testing. PLoS Comput Biol 13, e1005562 (2017)

17. R. Lyne, R. Smith, K. Rutherford, M. Wakeling, A. Varley, F. Guillier, H. Janssens, W. Ji, P. McLaren, P. North, D. Rana, T. Riley, J. Sullivan, X. Watkins, M. Woodbridge, K. Lilley, S. Russell, M. Ashburner, K. Mizuguchi, G. Micklem, FlyMine: an integrated database for Drosophila and Anopheles genomics. Genome Biol 8, R129 (2007)

